# Monocytes reprogrammed by 4-PBA potently contribute to the resolution of inflammation and atherosclerosis

**DOI:** 10.1101/2023.10.19.563200

**Authors:** Shuo Geng, Ran Lu, Yao Zhang, Yajun Wu, Ling Xie, Blake Caldwell, Kisha Pradhan, Ziyue Yi, Jacqueline Hou, Feng Xu, Xian Chen, Liwu Li

**Author notes:** Correspondence: Liwu Li, 970 Washington Street Virginia Tech, Blacksburg, VA 24061-0910.

## Abstract

**Background:** Chronic inflammation initiated by inflammatory monocytes underlies the pathogenesis of atherosclerosis. However, approaches that can effectively resolve chronic low-grade inflammation targeting monocytes are not readily available. The small chemical compound 4-phenylbutyric acid (4-PBA) exhibits broad anti-inflammatory effects in reducing atherosclerosis. Selective delivery of 4-PBA reprogrammed monocytes may hold novel potential in providing targeted and precision therapeutics for the treatment of atherosclerosis.

**Methods:** Systems analyses integrating single-cell RNA-sequencing and complementary immunological approaches characterized key resolving characteristics as well as defining markers of reprogrammed monocytes trained by 4-PBA. Molecular mechanisms responsible for monocyte reprogramming was assessed by integrated biochemical and genetic approaches. The inter-cellular propagation of homeostasis resolution was evaluated by co-culture assays with donor monocytes trained by 4-PBA and recipient naïve monocytes. The *in vivo* effects of monocyte resolution and atherosclerosis prevention by 4-PBA were assessed with the high fat diet-fed *ApoE^-/-^* mouse model with *i.p.* 4-PBA administration. Furthermore, the selective efficacy of 4-PBA trained monocytes were examined by *i.v.* transfusion of *ex vivo* trained monocytes by 4-PBA into recipient high fat diet-fed *ApoE^-/-^* mice.

**Results:** In this study, we found that monocytes can be potently reprogrammed by 4-PBA into an immune-resolving state characterized by reduced adhesion and enhanced expression of anti-inflammatory mediator CD24. Mechanistically, 4-PBA reduced the expression of ICAM-1 via reducing peroxisome stress and attenuating SYK-mTOR signaling. Concurrently, 4-PBA enhanced the expression of resolving mediator CD24 through promoting PPARγ neddylation mediated by TOLLIP. 4-PBA trained monocytes can effectively propagate anti-inflammation activity to neighboring monocytes through CD24. Our data further demonstrated that 4-PBA trained monocytes effectively reduce atherosclerosis pathogenesis when administered *in vivo*.

**Conclusion:** Our study describes a robust and effective approach to generate resolving monocytes, characterizes novel mechanisms for targeted monocyte reprogramming, and offers a precision-therapeutics for atherosclerosis based on delivering reprogrammed resolving monocytes.

## Introduction

Atherosclerosis is a complex chronic inflammatory disease culminating in the build-up of lipid-laden plaques within arterial vessels. Existing translational efforts are largely focused on developing chemical or biologic-based therapies that target metabolic and/or inflammatory processes ^1–3^. However, these approaches suffer from inherent drawbacks such as limited delivery precision into inflamed tissues and unintended side effects. As an alternative approach, mobilizing immune cells naturally equipped with effective physiological tropism into inflamed tissues may hold powerful therapeutic potentials in circumventing the drawbacks associated with molecule-based therapies.

To fully harness immune cell-based therapies, fundamental studies navigating the complex dynamics of atherosclerosis-associated immune cells are urgently needed. Emerging studies have identified monocytes as one of the most relevant immune cells involved in both the progression and resolution of atherosclerosis ^4–7^. The initial polarization of inflammatory monocyte subsets (e.g. Ly6C^hi^ murine monocytes or CD14^+^; CD16^+^ intermediate human monocytes) may serve as a key trigger for tissue infiltration, adhesion, and subsequent foamy macrophage formation ^8,9^. Strategies that can prevent the initial expansion of inflammatory monocytes would assist in the attenuation of atherosclerosis pathogenesis. Intriguingly, *in vivo* studies have also revealed that certain subsets of Ly6C^hi^ monocytes with limited proliferation potential can serve as precursors for anti-inflammatory resolving monocytes beneficial for atherosclerosis regression ^10–12^. Our recent *in vitro* studies with scRNA-seq pseudotime analyses further validated these *in vivo* observations by revealing the presence of distinct subsets of proliferative monocytes adopting either inflammatory or resolving properties ^12–14^. Following de-differentiation into a proliferative Ly6C^hi^ high state ^6,15^, monocytes bifurcate into either an inflamed state with the sustained stimulation of a low-grade inflammatory signal ^6^ or remain in an anti-inflammatory proliferative state ^15^. A clear characterization and derivation of resolving monocytes may facilitate future therapeutic development of monocyte-based precision therapies against atherosclerosis.

We recently reported that monocytes trained with small chemical compound 4-phenylbutyric acid (4-PBA) are arrested at the proliferative resolving state ^15,16^. 4-PBA is a potent peroxisome activator and can induce the expression of a PPAR-mediated peroxisome synthesis program as well as other anti-inflammatory mediators ^16–18^. Independent studies reported that the administration of 4-PBA into whole animals can reduce the progression of experimental atherosclerosis ^19–21^. In order to avoid the potential toxic effects of the systemic 4-PBA administration, we tested the approach of employing 4-PBA trained monocytes in delivering targeted therapeutics against atherosclerosis. Through integrated *in vitro* and *in vivo* studies, we defined key resolving characteristics of 4-PBA programmed monocytes, their underlying molecular mechanisms, and their therapeutic potential in attenuating atherosclerosis.

## Methods

### scRNA-seq data re-analysis

We performed re-analyses of publically deposited scRNAseq data sets processed as we described previously comparing naïve murine monocytes with monocytes trained by 4-PBA ^15^. Differentially expressed genes were analyzed using the non-parametric Wilcoxon rank sum test with the R package as we described ^15^. Genes demonstrating significant difference in expression levels were quantified with the Seurat R package and represented in the volcano plot. The dot plot was generated with the dot size representing the percentage of cell within the cell population expressing the gene of interest and the color intensity representing the normalized gene expression levels as we previously described ^6^. RNA-seq data have been uploaded to the NCBI Gene Expression Omnibus and are accessible under accession number GSE160450.

### Experimental animals

WT C57BL/6 mice, *ApoE*^−/−^ mice, and B6 SJL mice were purchased from the Jackson Laboratory. *Tram*^−/−^ mouse colony on C57BL/6 background was provided by Dr. Holger Eltzschig (University of Texas Houston Medical School, USA). *Tollip*^−/−^ mouse colony on C57BL/6 background was provided by Dr. Jürg Tschopp (University of Lausanne, Switzerland). *Cd24*^−/−^ mouse colony on C57BL/6 background was provided by Dr. Yang Liu (University of Maryland, USA). The mice were bred and maintained under pathogen–free conditions in the animal facility at Virginia Tech. All animal procedures were in accordance with the U.S. National Institutes of Health Guide for the Care and Use of Laboratory Animals and approved by the Intuitional Animal Care and Use Committee of Virginia Tech. Both male and female mice between 7 and 10 weeks of age were used for experiments, and no sex-specific effects were observed. Animals that showed health concerns unrelated to the experimental conditions (for example, fight wounds and dermatitis) were excluded.

### *In vitro* priming of mouse monocytes and flow cytometry analyses

Mouse BMM cultures were prepared as described before ^15,22^. BM cells isolated from WT C57BL/6 mice and *Tollip*^−/−^ mice were cultured in complete RPMI 1640 medium supplemented with 10 ng/mL M-CSF (Peprotech) in the presence of 1 mM 4-PBA (MilliporeSigma), 10 µg/mL oxLDL (Kalen Biomedical), 10 µg/mL cholesterol (MilliporeSigma), or PBS. Fresh 4-PBA, oxLDL, cholesterol or PBS was added to the cell cultures every 2 days. After 5 days, BMMs were harvested, filtered through 70 μm cell strainer, and incubated with anti-CD16/CD32 antibodies (1:100 dilution, BD Biosciences, no. 553141) to block Fc receptors. To determine surface expression of ICAM-1 and CD24, the BMMs were stained with fluorochrome-conjugated anti– CD11b (1:200 dilution, BioLegend, no. 101226), anti–Ly6C (1:200 dilution, BioLegend, no. 128018), anti–ICAM-1 (1:200 dilution, BioLegend, no. 116120), and anti–CD24 (1:200 dilution, BioLegend, no. 138504 or no. 101806) antibodies. Propidium iodide (MilliporeSigma) was added before flow cytometry to label dead cells. To detect intracellular mTOR expression, BMMs were first stained with fluorochrome-conjugated anti–CD11b (1:200 dilution, BioLegend, no. 101208), and anti–Ly6C (1:200 dilution, BioLegend, no. 128016) antibodies, fixed and permeabilized using Cyto-Fast™ Fix/Perm Buffer Set (BioLegend), and then stained with fluorochrome-conjugated anti–mTOR antibody (1:200 dilution, Cell Signaling Technology, no. 5043S). The samples were examined using FACSCanto II (BD Biosciences), and data were analyzed using FlowJo (Tree Star).

### HFD feeding, 4-PBA injection, and adoptive transfer of *in vitro* primed monocytes

Age- and gender-matched *ApoE*^−/−^ mice were fed with HFD (Harlan Teklad 94059) for 4 weeks to induce atherosclerosis development, and then the mice were intraperitoneally injected with PBS or 4-PBA (5 mg/kg body weight) every 3 days for 4 weeks. The mice were continuously fed with HFD during the regimen. One day after the last injection, the mice were euthanized, and tissues were harvested for subsequent analyses. Adoptive transfer of *in vitro* primed monocytes was conducted as reported previously ^6,22^. BM cells isolated from *ApoE*^−/−^ mice were cultured in complete RPMI 1640 medium supplemented with M-CSF (10 ng/mL) in the presence of 4-PBA (1 mM), or PBS for 5 days as described above. Age- and gender-matched recipient *ApoE*^−/−^ mice were first fed with HFD for 4 weeks to induce atherosclerosis development, and were then transfused with 3 × 10^6^ in vitro primed BMMs through intravenous injection once a week for a total of 4 weeks. In the experiments to test the influence of transferred monocytes on host cells, the in vitro primed monocytes were firstly labeled with CFSE (3 µM) before the adoptive transfer. The recipient mice were continuously maintained on HFD. One week after the last cell transfer, the mice were euthanized, and tissues were harvested for subsequent analyses.

### Histological analyses of atherosclerotic lesions

Histological analyses of atherosclerotic lesions were performed following established protocols ^6,22^. Briefly, 10 μm-thick sections of the proximal aorta, which had been freshly frozen and embedded in optimal cutting temperature (OCT) compound, were fixed in 4% neutral buffered formalin and subsequently subjected to Hematoxylin and Eosin (H&E) staining. Oil Red O staining was performed using a kit (Newcomer Supply), and collagen staining was performed using a Picrosirius Red Stain Kit (Polysciences) according to the manufacturers’ instructions. The samples were observed under a light microscope. The percentages of total lesion area, lipid deposition, and collagen composition were quantified.

### Determination of plasma lipids

Plasma samples were collected from the mice treated as described above. Total and free cholesterol levels were quantified with a kit purchased from MilliporeSigma, and triglyceride level was quantified with a kit purchased from BioVision. All assays were performed according to the manufacturers’ instructions.

### Flow cytometry analyses of monocyte phenotype *in vivo*

*ApoE*^−/−^ mice were fed with HFD and injected with 4-PBA as described above. Peripheral blood, BM, spleen, and aorta were harvested, and single-cell suspensions were prepared for flow cytometry as reported previously (ref). Briefly, blood, BM, and spleen samples were disassociated with mechanical processes followed by lysis of red blood cells with ACK buffer (Thermo Fisher Scientific). Aorta samples were cut into small pieces and then transferred into an enzyme cocktail containing 450 U/mL collagenase type I (Worthington), 250 U/mL collagenase type XI (MilliporeSigma), 120 U/mL hyaluronidase (MilliporeSigma), and 120 U/mL DNAse (MilliporeSigma), and 20 mM HEPES (MilliporeSigma). The samples were incubated at 37°C in an automated tissue dissociator (Miltenyi Biotec) for 60 minutes, and red blood cells were lysed with ACK buffer (Thermo Fisher Scientific). The processed samples of all tissues were filtered through 70 μm cell strainers to obtain single-cell suspensions. The samples were incubated with anti-CD16/-CD32 antibodies (1:100 dilution, BD Biosciences, no. 553141) to block Fc receptors followed by staining with fluorochrome-conjugated anti–CD11b (1:200 dilution, BioLegend, no. 101226), anti–Ly6C (1:200 dilution, BioLegend, no. 128018), anti–Ly6G (1:200 dilution, BioLegend, no. 127616, 127608 or 127616), anti–ICAM-1 (1:200 dilution, BioLegend, no. 116120 or 116108), and anti–CD24 (1:200 dilution, BioLegend, no. 101806 or 101808) antibodies. Propidium iodide (MilliporeSigma) was added before flow cytometry to label dead cells. The samples were then examined using FACSCanto II (BD Biosciences), and data were analyzed using FlowJo (Tree Star).

### Western blotting

BMMs from WT C57BL/6 and *Tram*^−/−^ mice were treated with 4-PBA (1 mM), oxLDL (10 µg/mL), or PBS for 5 days as described above. Total protein was extracted with RIPA buffer (Thermo Fisher Scientific) containing a protease inhibitor cocktail (MilliporeSigma) and a phosphatase inhibitor cocktail (MilliporeSigma). Protein samples were subjected to SDS-PAGE and transferred to a polyvinylidene difluoride membrane. The membrane was incubated with blocker (Bio-Rad) at room temperature for 1 hour and then incubated overnight at 4°C with primary anti–SYK (1:1000 dilution, Cell Signaling Technology, no. 2712S), anti–p-p38 (1:1000 dilution, Cell Signaling Technology, no. 4511S), or anti–β-actin antibody (1:1000 dilution, Cell Signaling Technology, no. 5125), followed by incubation with HRP-conjugated anti–rabbit IgG (1:1000 dilution, Cell Signaling Technology, no. 7074) for 1 hour at room temperature. Blots were developed by an enhanced chemiluminescence (ECL) detection kit (Thermo Fisher Scientific) and detected with Amersham Imager 600 (GE HealthCare).

### Confocal microscopy

To detect lysosomal distribution of mTOR and SYK, WT BMMs were treated with 4-PBA (1 mM), oxLDL (10 µg/mL), or PBS for 5 days as described above. The cells were rinsed with PBS, fixed with 4% paraformaldehyde, and permeabilized with 0.2% Triton X-100. After blocking with 10% goat serum, the samples were stained with primary rabbit anti–mouse PMP70 antibody (1:1000 dilution), followed by staining with Alexa Fluor 488 goat anti–rabbit secondary antibody (1:1000 dilution). Paraformaldehyde, Triton X-100, goat serum, rabbit anti–mouse PMP70 antibody, and Alexa Fluor 488 goat anti–rabbit secondary antibody were all supplied in the SelectFX Alexa Fluor 488 peroxisome labeling kit (Thermo Fisher Scientific). After washing with PBS, the cells were stained with CoraLite^®^594 conjugated anti–mTOR antibody (1:300 dilution; Proteintech, no. CL594-66888) or PE conjugated anti–SYK antibody (1:500 dilution; BioLegend, no. 646004). After washing with PBS, the samples were mounted with antifade mountant (Invitrogen). To detect the distribution of TRAM, WT BMMs were treated with 4-PBA (1 mM), oxLDL (10 µg/mL), cholesterol (10 µg/mL) or PBS for 5 days as described above. The cells were rinsed with PBS, fixed with 4% paraformaldehyde, and permeabilized with 0.2% Triton X-100. After blocking with 10% goat serum, the samples were stained with biotin conjugated anti–mouse TRAM antibody (1:500 dilution, Biorbyt, no. orb452158), followed by staining with Alexa Fluor 488 conjugated streptavidin (1:1000 dilution, Thermo Fisher Scientific, no. S32354) After washing with PBS, the samples were mounted with antifade mountant (Invitrogen). All the samples were observed under an LSM 900 confocal microscope (ZEISS). Images were processed with ZEN lite (ZEISS).

### Immunoprecipitation

Immunoprecipitation was conducted using Pierce™ Co-Immunoprecipitation Kit (Thermo Fisher Scientific) according to the manufacturer’s instructions. To screen the proteins associated with Tollip, antibodies against Tollip (NOVUS, no. H00054472-M01) were conjugated to the resin supplied in the kit. WT BMMs were cultured for 5 days, and cell lysate was isolated and then incubated with antibody-conjugated resin overnight at 4°C. Immunoprecipitated products were eluted and subjected to mass spectrometry analysis. To determine the neddylation of PPARγ, antibodies against NEDD8 (Thermo Fisher Scientific, no. MA5-32552) were conjugated to the resin. BMMs from WT and *Tollip*^−/−^ mice were treated with 4-PBA (1 mM) or PBS for 5 days as described above. Equal amounts of cell lysate samples were incubated with antibody-conjugated resin overnight at 4°C. Immunoprecipitated products were eluted and subjected to Western blotting with primary anti-PPARγ antibody (1:1000 dilution, Thermo Fisher Scientific, no. PA3-821A).

### Mass spectrometry analysis

Tollip immunoprecipitates were eluted with 8 M urea. The eluate was reduced with 5 mM Dithiothreitol (DTT) and alkylated with 15 mM Iodoacetamide (IAA) followed by overnight trypsin digestion. The peptides were cleaned with home-made C18 stage-tips. The clean peptides were dissolved in 0.1% formic acid and analyzed on a Q-Exactive Orbitrap mass spectrometer coupled with an Easy nanoLC 1000 (Thermo Fisher Scientific, San Jose, CA). Peptides were loaded on to a 15 cm C18 RP analytical column (75 μm inner diameter, 2 μm, 100-Å particle, Acclaim Pepmap RSLC, Thermo-Fisher). Analytical separation of all peptides was achieved with 60-min gradient. A linear gradient of 2 to 30% buffer B over 40 min and 30% to 80% buffer B over 5 min was executed at a 300 nl/min flow rate followed 15-min wash with 100%B, where buffer A was aqueous 0.1% formic acid, and buffer B was 100% acetonitrile and 0.1% formic acid. LC-MS experiments were performed in a data-dependent mode with full MS at a resolution of 70,000 at m/z 200 followed by high energy collision-activated dissociation-MS/MS of the top 20 most intense ions with a resolution of 17,500 at m/z 200. High energy collision-activated dissociation-MS/MS was used to dissociate peptides at a normalized collision energy of 27 eV in the presence of nitrogen bath gas atoms. Dynamic exclusion was 15.0 seconds. Mass spectra were processed, and peptide identification was performed using the MaxQuant software version 1.5.3.17 (Max Planck Institute, Germany). Protein database searches were performed against the UniProt human protein sequence database. A false discovery rate (FDR) for peptide-spectrum match (PSM) was set at 1% and FDR for protein assignment was set at 5%. Search parameters included up to two missed cleavages at Lys/Arg on the sequence, oxidation of methionine, and protein N-terminal acetylation as a dynamic modification. Carbamidomethylation of cysteine residues was considered as a static modification. Peptide identifications are reported by filtering of reverse and contaminant entries and assigning to their leading razor protein.

### Co-culture assay

BMMs from B6 SJL mice (CD45.1^+^) were used as recipient cells; BMMs from WT C57BL/6 mice, *Cd24*^−/−^ mice, and *Tollip*^−/−^ mice (CD45.2^+^) were used as donor cells. Co-culture assay was performed using previously established protocol with modifications ^6^. Briefly, donor monocytes were treated with 4-PBA (1 mM) or PBS for 5 days, and an equal number of primed donor cells were added to recipient monocytes that had been treated with oxLDL (10 µg/mL) for 3 days. The donor and recipient cells were subsequently co-cultured for additional 2 days in the presence of M-CSF (10 ng/mL). The cells were harvested and stained with anti–CD45.1 (1:200 dilution, BioLegend, no. 110730), anti–CD24 (1:200 dilution, BioLegend, no. 101806), and anti–ICAM-1 (1:200 dilution, BioLegend, no. 116108) antibodies. The surface phenotype of CD45.1^+^ recipient BMMs was determined with flow cytometry.

### ELISA

For the detection of CCL5 produced by monocytes *in vitro*, WT and *Tram*^−/−^ BMMs were treated with 4-PBA (1 mM), oxLDL (10 µg/mL), cholesterol (10 µg/mL) or PBS for 5 days as described above, and supernatant was collected. For the detection of CCL5 level *in vivo*, plasma samples were collected from the atherosclerotic mice that received monocyte transfusion as described above. The concentration of CCL5 in the supernatant and plasma samples was assessed with an ELISA kit (R&D Systems).

### Adhesion assay

Adhesiveness of monocytes was measured with Vybrant™ Cell Adhesion Assay Kit (Thermo Fisher Scientific) according to published method ^23^. WT BMMs were treated with 4-PBA (1 mM) or PBS for 5 days. The cells were harvested and labeled with calcein AM (5 µM) at 37°C for 30 minutes. An equal number of calcein-labeled monocytes were added to each well of a 96-well plate followed by incubation at 37°C for 2 hours. Non-adherent cells were removed by careful washing, and fluorescence from adherent cells was measured.

### Detection of monocyte migratory capacity

WT BMMs were treated with 4-PBA (1 mM) or PBS as described above. After 5 days, the monocytes were harvested, and the migratory capacity was examined with the method reported previously ^24^. Monocytes were suspended in complete RPMI containing M-CSF (10 ng/mL) and loaded into polystyrene crosshatched nanofiber scaffolds. The samples were placed in an incubation chamber set at 37°C with 5% CO_2_ and observed under Axio Observer Z1 microscope with digitally controlled three-axis stage (ZEISS). The movement of monocytes during 10 hours was recorded with 5-minute intervals between images. Instantaneous velocity of monocytes was analyzed with ImageJ.

### Assessment of monocyte proliferation *in vivo*

Age- and gender-matched *ApoE*^−/−^ mice were fed with HFD and intraperitoneally injected with PBS or 4-PBA (5 mg/kg body weight) as described above. Proliferation of monocytes was analyzed with In Vivo EdU Flow Cytometry 50 Kit 488 (MilliporeSigma) following the manufacturers’ instructions. Briefly, EdU (50 mg/kg body weight) was intraperitoneally injected to the mice 4 hours before euthanasia, and BM and spleen were harvested to prepare single cell suspensions. The cells were first stained with anti–CD11b (1:200 dilution, BioLegend, no. 101212), anti–Ly6C (1:200 dilution, BioLegend, no. 128026), and anti–Ly6G (1:200 dilution, BioLegend, no. 127616) antibodies against surface antigens. After fixation and permeabilization, the cells were incubated with Click-iT^®^ reaction cocktail supplied in the kit. EdU^+^ population within Ly6C^hi^ monocytes was examined by flow cytometry.

### Seahorse assay

WT BMMs were seeded the Seahorse Bioscience XF24 (Agilent) cell culture plates and treated with oxLDL (10 µg/mL), low dose LPS (100 pg/mL) or PBS for 5 days. Cell Mito Stress Test was conducted by sequentially exposing the cells to 0.5 μM oligomycin, 0.6 μM carbonyl cyanide-4-(trifluoromethoxy) phenylhydrazone (FCCP), and 0.1 μM rotenone/antimycin A, in accordance with the manufacturer’s instructions.

### Genomic methylation analysis

WT BMMs were treated with cholesterol (10 µg/mL), low dose LPS (100 pg/mL), high dose LPS (1 µg/Ml) or PBS for 5 days. Genomic DNA was prepared from cultured monocytes using a DNeasy Blood & Tissue Kit (QIAGEN), bisulfite-treated using an EpiTect Bisulfite Kit (QIAGEN), and processed and hybridized to individual array wells of an Infinium Mouse Methylation BeadChip (Illumina), as previously described ^25^. All arrays were processed and run at the Children’s Hospital of Philadelphia Center for Applied Genomics. Raw Infinium IDAT files were processed into corrected beta values using the openSesame pipeline (1.14.2; default parameters), and differentially methylated regions (DMRs) were identified using the sesame DML function ^26^. PBS- and LPS-treated cultured bone marrow monocyte array values were previously published, but were collected and sequenced at the same time as low dose LPS- and cholesterol-stimulated samples ^25^. Infinium BeadChip probe annotations were obtained using the sesame KYCG function ^26^. Principal components analysis was performed using the R stats package. Methylation profiling data have been uploaded to the NCBI Gene Expression Omnibus and are accessible under accession number GSE243358.

### *In vitro* priming of human monocytes and flow cytometry analyses

Peripheral blood collected from healthy individuals was purchased from Research Blood Components, LLC (Boston, MA). Peripheral blood mononuclear cells (PBMCs) were isolated using Ficoll-Paque^TM^ (Cytiva), and then cultured with M-CSF (100 ng ml−1) in the presence of 4-PBA (1 mM) or PBS. After 2 days, cells were harvested, blocked with TruStain FcX™ (BioLegend) and stained with fluorochrome-conjugated anti–CD14 (1:50 dilution, BioLegend, no. 325618), anti–CD16 (1:50 dilution, BioLegend, no. 360710), anti–ICAM-1 (1:50 dilution, BioLegend, no. 353114), or anti–CD24 (1:50 dilution, BioLegend, no. 311106) antibodies. Propidium iodide (MilliporeSigma) was added before flow cytometry to label dead cells. The samples were then examined using FACSCanto II, and data were analyzed using FlowJo.

### Statistics

Statistical analyses were performed using Prism software (GraphPad). All data are expressed as means ± SEM, and the sample number for each data set is provided in figure legends. Comparisons between 2 groups were performed by using 2-tailed Student’s t test, and comparisons among multiple groups were carried out with 1-way ANOVA. *P* < 0.05 was considered statistically significant.

## Results

### Monocytes trained with 4-PBA exhibit potent anti-inflammatory resolving characteristics

Administration of 4-PBA was independently shown to be beneficial in reducing atherosclerosis ^19–21^, indicating that 4-PBA may serve as a promising atheroprotective agent. Our *in vitro* studies reveal that monocytes treated with 4-PBA are arrested in an anti-inflammatory state, suggesting monocytes may be responsible for atheroprotective benefits of 4-PBA in animal models. To better define and harness the therapeutic potential of resolving monocytes programed by 4-PBA, we re-analyzed the gene expression profile of monocytes treated with 4-PBA and monitored for key signatures indicative of monocyte differentiation and activation. As shown in Figure 1, 4-PBA trained monocytes express monocytic markers such as *Cd34* and have drastically reduced expression of mature macrophage markers such as *Adgre1* (F4/80), consistent with morphological observation (Figure S1A) demonstrating the monocytic nature of 4-PBA trained monocytes as compared to control monocyte cultures. As a consequence, 4-PBA trained monocytes exhibit significantly reduced adhesion capacity and elevated migratory potential (Figure S1B, Video S1, and Video S2). 4-PBA trained monocytes also express higher levels of genes representative of elevated proliferative potentials such as *Hmmr*, *Mki67* and *Stmn1* (Figure 1A-C). We next examined pro- and anti-inflammatory gene signatures, and observed lower levels of adhesion molecules such as ICAM-1 and drastically elevated anti-inflammatory mediators such as CD24. ICAM-1 plays a pivotal role as a pro-inflammatory biomarker that facilitates monocyte adhesion to aortic endothelial cells and contributes to the development of atherosclerotic plaques ^27,28^. The expression of CD24 is found on the surface of various hematopoietic cell populations, and thus CD24 expression is higher in progenitor cells and less differentiated cells as compared to terminally differentiated cells ^29^. CD24 has also been well-documented to interact with SIGLEC10 to suppress inflammatory responses of innate immune cells to infection, sepsis, and chronic inflammatory diseases ^30^. To validate the expression of ICAM-1 and CD24 in 4-PBA-treated cells at the protein levels, we first employed our previously established murine monocyte culture system. ^16,22^ In line with our scRNA-seq results, flow cytometry demonstrated that treatment with 4-PBA (1 mM) for 5 days significantly reduced ICAM-1 expression and elevated CD24 expression on murine bone marrow-derived monocytes (BMMs) compared to PBS control cells (Figure 1D). To validate the translational relevance of these findings, we tested whether 4-PBA can elicit similar pro-resolving characteristics in human primary monocytes. We cultured human peripheral blood mononuclear cells (PBMCs) with either PBS or 4-PBA and observed a significant reduction of ICAM-1 and elevation of CD24 in 4-PBA-treated cells (Figure 1E). Our data reveal that 4-PBA programmed monocytes adopt key resolving signatures of less-differentiated monocytes with potential atheroprotective effects.

**Figure 1.**
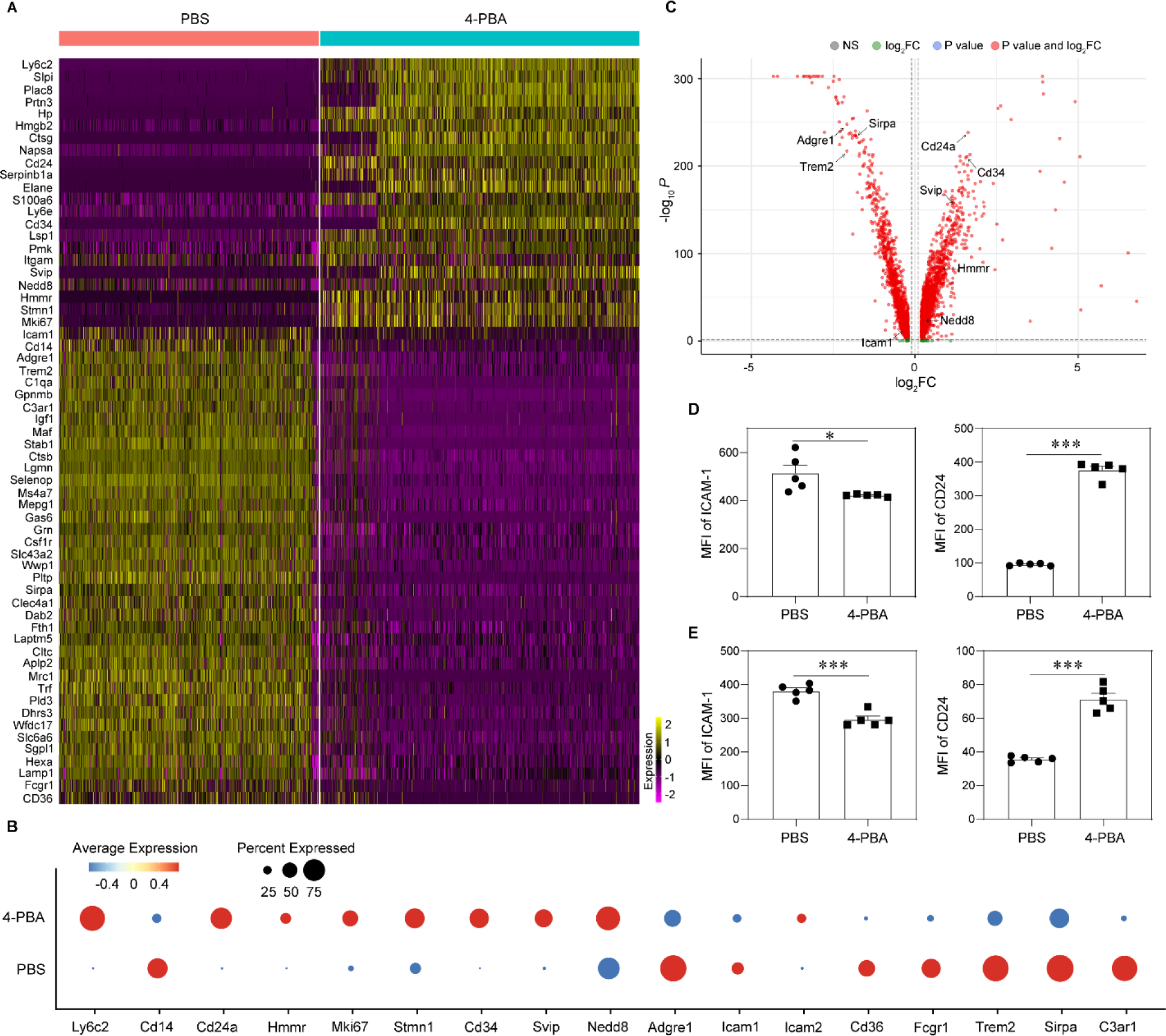
Treatment with 4-PBA induces anti-inflammatory resolving characteristics of monocytes. **A-D,** BMMs from WT C57 BL/6 mice were cultured with M-CSF (10 ng/mL) in the presence 4-PBA (1 mM) or PBS for 5 days. scRNA-Seq was performed and data sets processed to comparing naïve murine monocytes with monocytes trained by 4-PBA. **A,** Heatmaps demonstrating representative genes differentially expressed in different clusters of monocytes challenged with 4-PBA. **B,** Dot plot comparison of representative genes differentially expressed between PBS-versus 4-PBA-primed monocytes. **C,** Volcano plot of data as log_2_ fold change versus the −log_10_ of the adjusted P value. Genes with significantly different are highlighted as red dots. **D,** Surface expression of ICAM-1 and CD24 on CD11b^+^ monocytes was analyzed by flow cytometry. **E,** PBMCs isolated from healthy human individuals were cultured with M-CSF (100 ng/mL) in the presence 4-PBA (1 mM) or PBS for 2 days. Surface expression of ICAM-1 and CD24 on total monocytes (including CD14^hi^, CD16^hi^ and intermediate monocytes) was analyzed by flow cytometry. Data in **D** and **E** (n = 5 for each group) are representative of three independent experiments. Error bars represent means ± SEM. \**P* < 0.05, \*\*\**P* < 0.001; Student’s 2-tailed t test.

### 4-PBA injection reprograms monocytes in vivo and reduces atherosclerosis

We next tested whether these key signatures of resolving monocytes can be recapitulated in mice treated with 4-PBA *in vivo*. We initially confirmed that administration of 4-PBA may alleviate atherosclerosis progression in our experimental mouse model. Whereas previous studies administrated 4-PBA through drinking water ^19,20^, we elected to treat high-fat diet (HFD)–fed *ApoE^−/−^* mice with 4-PBA via intraperitoneal (*i.p.*) injection (100 mg/kg body weight). *ApoE^−/−^* mice were HFD fed for 4 weeks to induce the development of atherosclerosis, and then received i.p. injections of 4-PBA every 3 days for an additional 4 weeks, during which the mice were continuously HFD fed. In comparison to the mice injected with vehicle control (PBS), those injected with 4-PBA exhibited significantly reduced size of atherosclerotic plaques, as evident from H&E staining (Figure 2A), as well as remarkably diminished lipid deposition in the plaques, as shown by Oil Red O staining (Figure 2B). Moreover, injection of 4-PBA significantly increased the collagen content within plaques, indicating an improvement of plaque stability (Figure 2C). The plasma levels of total cholesterol, free cholesterol, and triglyceride were also significantly reduced after 4-PBA administration (Figure 2D). Our data validated that long-term *i.p.* injection of 4-PBA drastically reduced the atherosclerotic burden in experimental mice.

**Figure 2.**
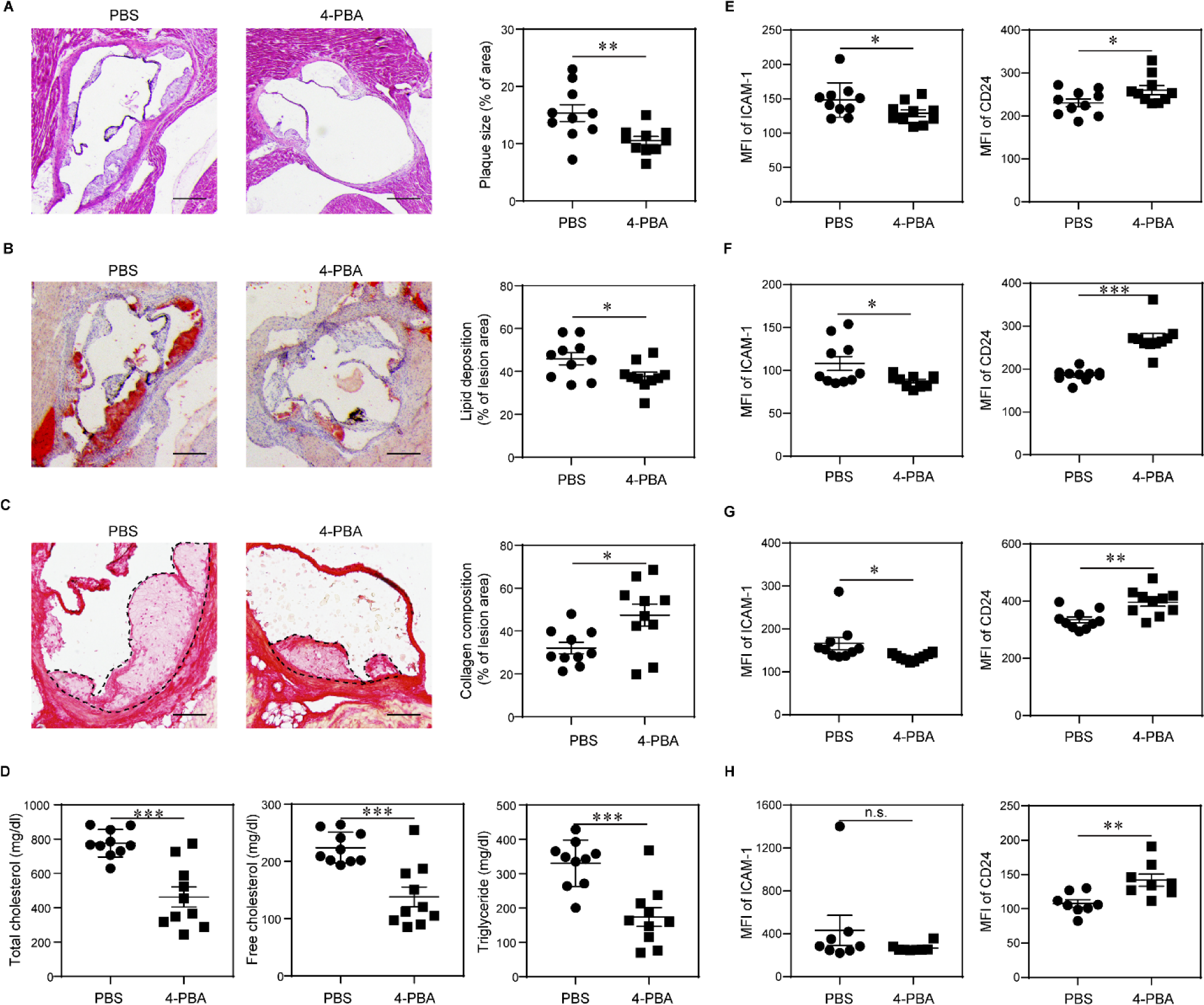
Administration of 4-PBA alleviates atherosclerotic pathogenesis. Male *ApoE*^−/−^ mice were fed with HFD for 4 weeks and intraperitoneally injected with 4-PBA (5 mg/kg body weight) or PBS every 3 days for additional 4 weeks. **A,** Representative images of H&E-stained atherosclerotic lesions and quantification of plaque size demonstrated as the percentage of lesion area within aortic root area. Scale bars, 300 μm. **B,** Representative images of Oil Red O–stained atherosclerotic plaques and quantification of lipid deposition within lesion area. Scale bars, 300 μm. **C,** Representative images of Picrosirius red–stained atherosclerotic plaques and quantification of collagen content within lesion area. Scale bars, 100 μm. **D,** Detection of total cholesterol, free cholesterol and triglyceride levels in the plasma. **E-F,** Surface expressions of ICAM-1 and CD24 on CD11b^+^ Ly6G^−^ Ly6C^hi^ monocytes in the peripheral blood (**E**), BM (**F**), spleen (**G**), and aorta (**H**) were examined by flow cytometry. Data are representative of two independent experiments (n = 10 for each group in **A-G**; n = 8 for each group in **H**). Error bars represent means ± SEM. \**P* < 0.05, \*\**P* < 0.01, \*\*\**P* < 0.001, n.s. no statistical significance; Student’s 2-tailed t test.

Consistent with prior independent studies, we also observed proliferating Ly6C^hi^ monocytes in atherosclerotic mice, as evidenced by *in vivo* Edu incorporation (Figure S2). Notably, administration of 4-PBA substantially increased the frequency of these proliferating monocytes in both the bone marrow and spleen (Figure S2). It is particularly noteworthy that relative to mice receiving vehicle controls, mice injected with 4-PBA exhibited significantly reduced levels of ICAM-1 and elevated levels of CD24 on Ly6C^hi^ and Ly6C^low^ monocytes harvested from the peripheral blood, bone marrow and, spleen, as well as aorta (Figure 2E-H and Figure S3). These data further substantiate that injection of 4-PBA prompts the polarization of monocytes to a resolving state in atherosclerotic mice, which potentially contributes to the amelioration of atherosclerosis pathogenesis.

### 4-PBA reduces monocyte adhesion by restoring pexophagy and reducing mTOR signaling

We further examined the molecular and cellular mechanisms responsible for the reprogramming of resolving monocytes by 4-PBA. Previous studies suggest that atherosclerotic stress factors such as oxLDL and/or cholesterol can initiate the inflammatory polarization of monocyte via activation of mTORC1, and that application of mTOR inhibitor rapamycin effectively reduces atherosclerosis progression ^31^. mTOR can be activated on the subcellular platforms of lysosomes or peroxisomes by reactive oxygen species (ROS) resulting from defective pexophagy ^32,33^. Consistent with previous reports, we noted that in contrast to high-dose LPS treatment, oxLDL or low-dose LPS did not compromise mitochondria function, but rather improved mitochondria respiration ^34^ (Figure S4). We therefore focused on the effects of oxLDL on inducing the dysfunction of peroxisomes. Flow cytometry analysis revealed a remarkable elevation of intracellular mTOR level in monocytes after treatment with oxLDL for 5 days (Figure 3A). Intriguingly, we observed that oxLDL drastically increased the subcellular localization of mTOR at PMP70^+^ peroxisomes (Figure 3B). Moreover, oxLDL also increased cellular levels and peroxisomal distribution of SRC kinase SYK (Figure 3C-D), which was shown to form a mutually activating positive feedback loop with mTOR with the support of subcellular ROS ^35^. These findings corroborate prior studies suggesting that peroxisomes may act as a pivotal platform for sustaining the inflammatory signaling cascade ^36^.

**Figure 3.**
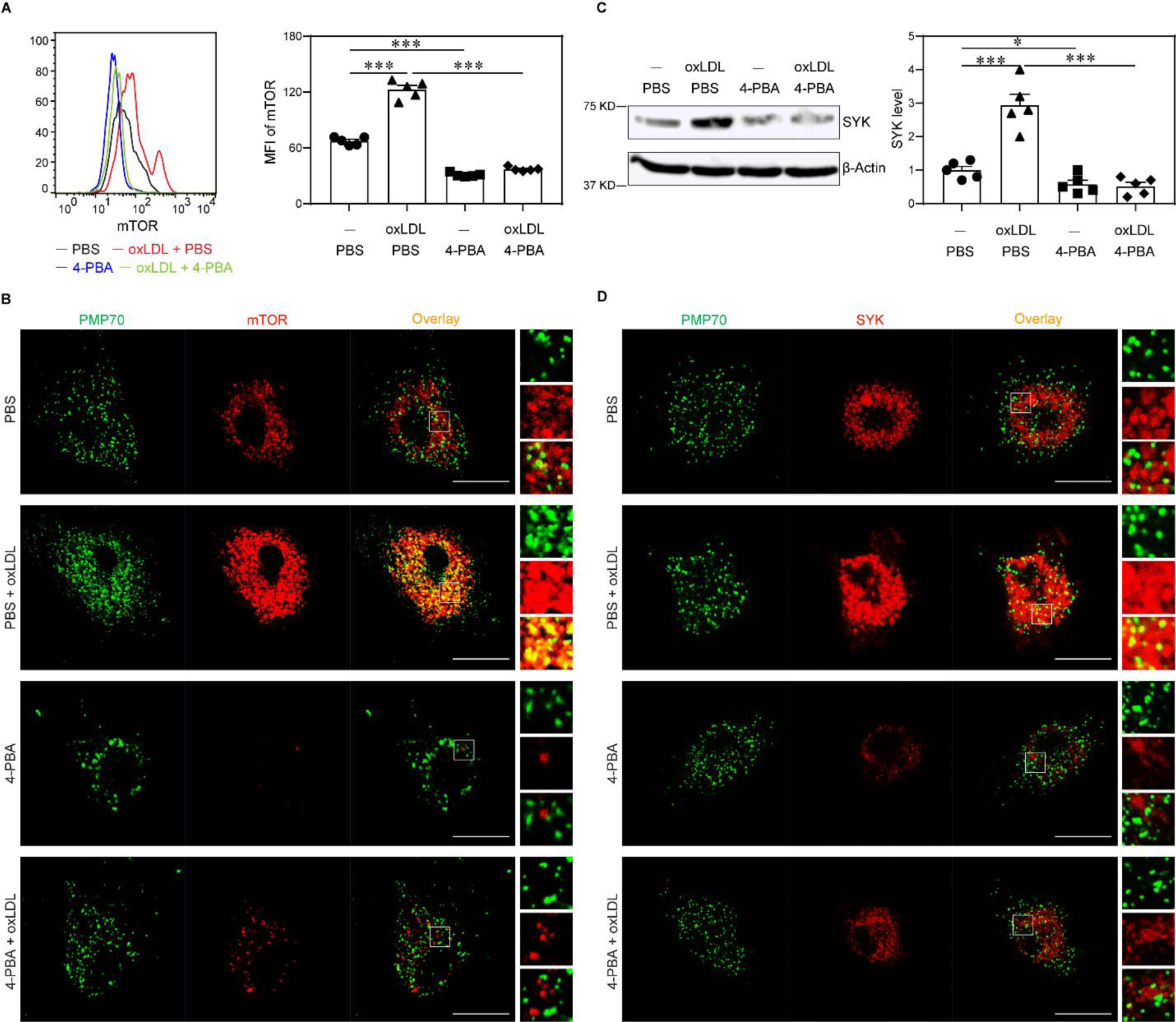
4-PBA inhibits mTOR signaling and restores peroxisome homeostasis in monocytes. BMMs from WT C57 BL/6 mice were cultured with M-CSF (10 ng/mL) in the presence of oxLDL (10 µg/mL), 4-PBA (1 mM) or PBS for 5 days. **A,** Representative histogram and quantification of mTOR level in CD11b^+^ Ly6C^hi^ monocytes as determined by flow cytometry. **B,** Monocytes were stained with anti–PMP70 and anti–mTOR antibodies, and the localization of PMP70^+^ peroxisomes and mTOR was examined by confocal microcopy. Scale bars: 10 μm. **C,** Protein level of SYK in BMMs was examined by Western blotting, and relative expression was normalized to β-actin. **D,** Monocytes were stained with anti–PMP70 and anti–SYK antibodies, and the localization of peroxisomes and SYK was examined by confocal microcopy. Scale bars: 10 μm. Data are representative of three independent experiments (n = 5 for each group). Error bars represent means ± SEM. \**P* < 0.05, \*\*\**P* < 0.001; one-way ANOVA.

We previously reported that membrane-associated adaptor TRAM (also known as TICAM-2) can transmit the signal of super-low dose LPS and is responsible for causing peroxisomal dysfunction ^6,37^, as well as inducing key low-grade inflammatory mediators ^6,38^. Given that oxLDL or free cholesterol can cause generic membrane stress ^39–41^, and that TRAM is one of the few innate membrane adaptors with lipid anchors to stressed membrane region ^42^, we next tested the hypothesis that TRAM serves as a general membrane stress sensor capable of mediating the inflammatory effects of oxLDL or free cholesterol in addition to low-dose LPS. As shown in Figure 4A, both oxLDL and free cholesterol promoted TRAM clustering on the cell membrane. We further examined cellular activation in wild type (WT) and TRAM-deficient monocytes challenged with oxLDL. We found that the potent activation of p38 as well as SYK by oxLDL is observed in WT but not *Tram*^-/-^ monocytes (Figure 4B-C). Functionally, oxLDL or free cholesterol induced ICAM-1 expression and CCL5 secretion in wild type monocytes, but not TRAM deficient monocytes (Figure 4D). Our data further confirm that oxLDL induces monocyte activation via a generic membrane-associated stress sensor TRAM.

**Figure 4.**
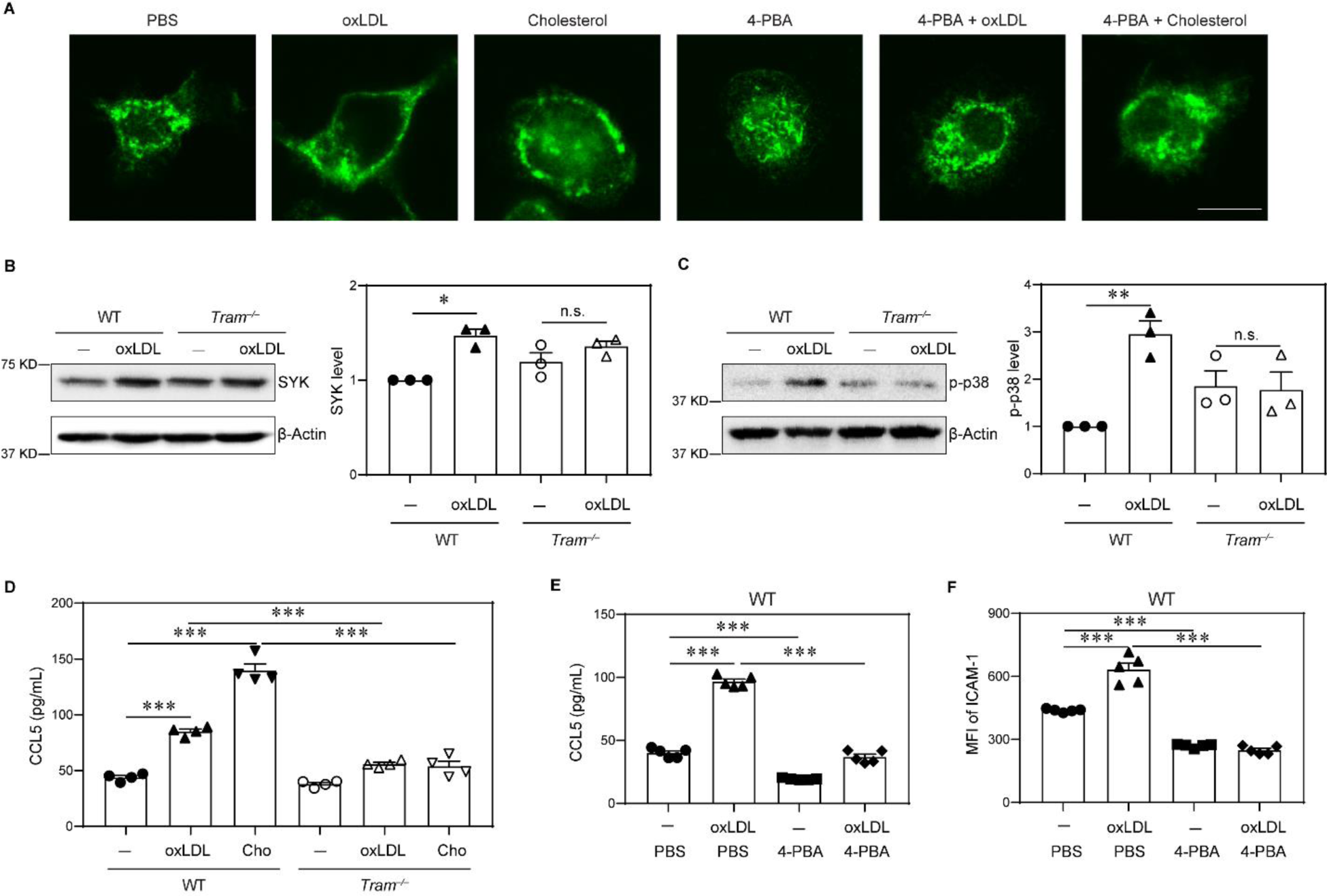
TRAM serves as a general membrane stress sensor mediating the inflammatory effects of lipids on monocytes. **A,** BMMs from WT C57 BL/6 mice were cultured with M-CSF (10 ng/mL) in the presence of oxLDL (10 µg/mL), cholesterol (10 µg/mL), 4-PBA (1 mM) or PBS for 5 days. The cells were stained with anti–TRAM antibody, and cellular distribution of TRAM was examined by confocal microcopy. Scale bars: 10 μm. **B**-**C,** BMMs from WT C57 BL/6 mice and *Tram*^−/−^ mice were cultured with M-CSF (10 ng/mL) in the presence of oxLDL (10 µg/mL) or PBS for 5 days. Protein level of SYK (**B**) and phosphorylation of p-38 (**C**) were examined by Western blotting, and relative expression was normalized to β-actin. **D,** BMMs from WT C57 BL/6 mice and *Tram*^−/−^ mice were cultured with M-CSF (10 ng/mL) in the presence of oxLDL (10 µg/mL), cholesterol (10 µg/mL) or PBS for 5 days. Production of CCL5 was determined by ELISA. **E-F,** BMMs from WT C57 BL/6 mice were cultured with M-CSF (10 ng/mL) in the presence of oxLDL (10 µg/mL), 4-PBA (1 mM) or PBS for 5 days. Production of CCL5 was determined by ELISA (**E**), and surface expression of ICAM-1 on CD11b^+^ monocytes was determined by flow cytometry (**F**). Data are representative of three independent experiments (n = 3 for each group in **B** and **C**; n = 4 for each group in **D**; n = 5 for each group in **E** and **F**). Error bars represent means ± SEM. \**P* < 0.05, \*\**P* < 0.01, \*\*\**P* < 0.001, n.s. no statistical significance; one-way ANOVA.

To further validate that low-dose LPS and cholesterol can similarly reprogram monocyte memory, we then examined genome wide methylation profiles of monocytes trained by varying dosages of LPS or cholesterol. Indeed, principle component analyses showed that exhausted monocytes caused by prolonged challenges with high dose LPS clustered into a distinct population in terms of their methylation profile. By contrast, monocytes programed by either low-dose LPS or cholesterol clustered together (Figure S5).

Based on these novel mechanistic findings, we next turned our attention back to the anti-inflammatory effects of 4-PBA. We tested whether 4-PBA treatment may diffuse the membrane clustering of TRAM via confocal microscopy. We observed that co-incubation of 4-PBA indeed diffused the membrane clustering of TRAM and caused a permissive cytosolic distribution of TRAM (Figure 4A). At the signaling level, we observed that 4-PBA treatment drastically reduced the cellular levels of SYK and mTOR, as well as the co-localization of mTOR and SYK with peroxisome (Figure 3). Functionally, we observed that 4-PBA significantly reduced the expression of ICAM-1 and CCL5 in oxLDL-treated cells (Figure 4E-F). Of note, CCL5 was recently implicated via a comprehensive bioinformatics study as the most relevant inflammatory mediator underlying human atherosclerosis ^43^. Collectively, our data reveal that 4-PBA can mitigate the pro-inflammatory memory of monocytes through the attenuation of TRAM-mediated subcellular inflammatory stress.

### 4-PBA promotes the expression of anti-inflammatory mediator CD24 through enhanced PPARγ activation

4-PBA not only reduced monocyte inflammation and the expression of adhesion molecule ICAM-1, but also significantly increased the expression of anti-inflammatory mediator CD24. To further characterize the molecular mechanism for elevated CD24 expression, we examined genes induced by 4-PBA in the scRNA-seq data set and noted the elevated expression of NEDD8 (Figure 1A-C). NEDD8-mediated PPARγ neddylation was shown to potently enhance PPARγ activation ^44^, which may be responsible for the elevated induction of CD24. We tested the status of PPARγ neddylation by co-immunoprecipitation and observed increased PPARγ neddylation in monocytes trained by 4-PBA (Figure 5A).

**Figure 5.**
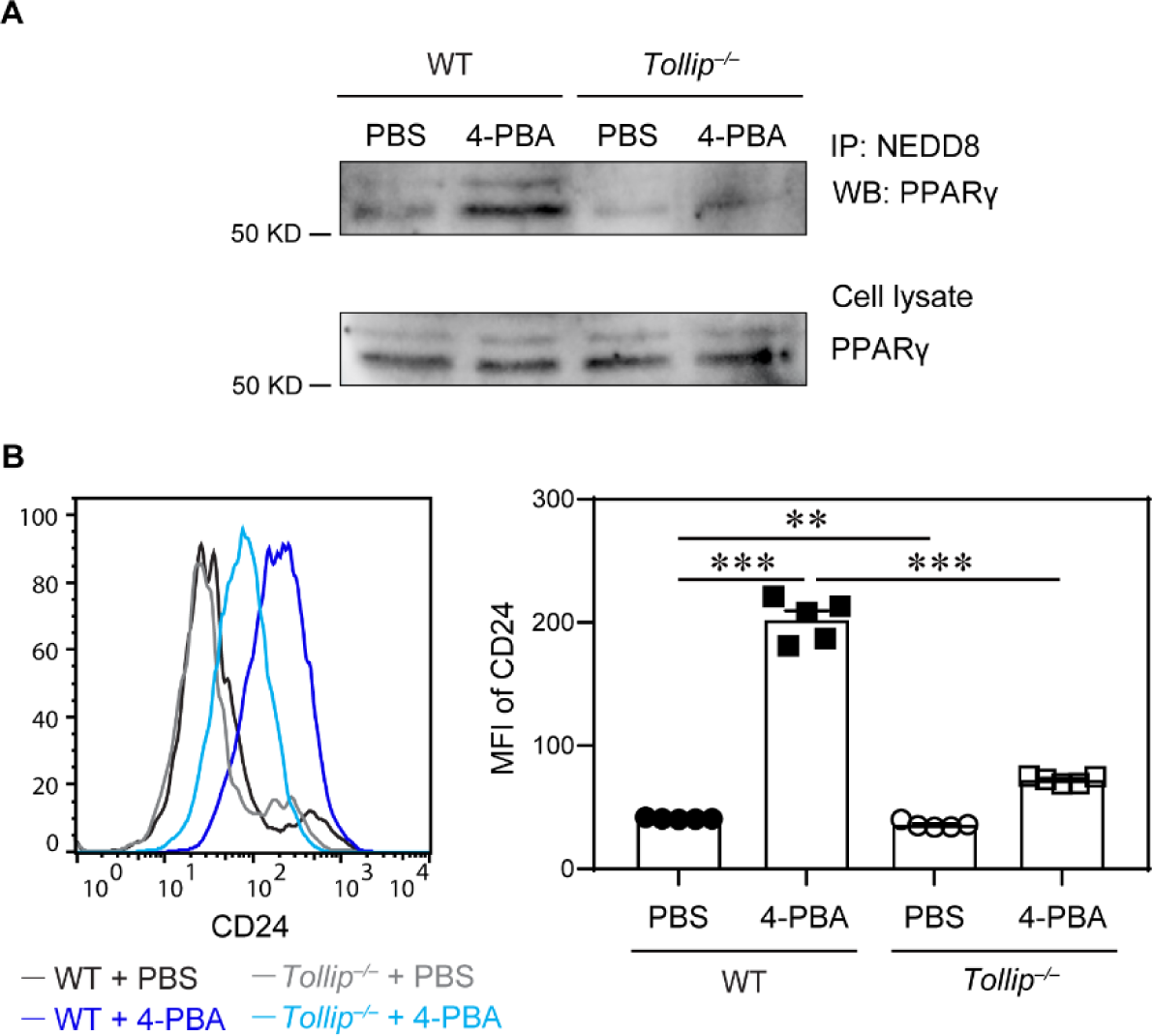
4-PBA promotes the expression of CD24 through enhanced PPARγ activation in a Tollip dependent manner. BMMs from WT C57 BL/6 mice and *Tollip*^−/−^ mice were cultured with M-CSF (10 ng/mL) in the presence of 4-PBA (1 mM) or PBS for 5 days. **A**, Cell lysate was isolated and subjected to immunoprecipitation with anti–NEDD8 antibodies conjugated to resin. The association of PPARγ with NEDD8 was examined by Western blotting. PPARγ in cell lysate was also examined by Western blotting. **B**, Surface expression of CD24 on CD11b^+^ Ly6G^−^ Ly6C^hi^ monocytes in the was examined by flow cytometry. Data are representative of three independent experiments (n = 5 for each group in **B**). Error bars represent means ± SEM. \*\**P* < 0.01, \*\*\**P* < 0.001; one-way ANOVA.

To clearly define molecular mechanisms responsible for increased PPARγ neddylation by 4-PBA, we next examined NEDD8-interacting molecule TOLLIP, discovered through an unbiased proteomic analysis of peptides co-precipitated with TOLLIP (Data Set 1). TOLLIP was also independently identified as an important mediator for subcellular organelle fusion and homeostasis ^45,46^. Furthermore, *Tollip* deletion results in the expansion of inflammatory monocytes and exacerbates atherosclerosis ^47^. We tested whether *Tollip* deficiency ablates the beneficial effects of 4-PBA on monocyte resolution. As shown in Fig. 5a, PPARγ neddylation was reduced in *Tollip*-deficient monocytes. Functionally, we observed that *Tollip* deletion significantly dampened the induction of CD24 on monocytes by 4-PBA (Figure 5B and Figure S6). Our data indicate that treatment with 4-PBA promotes PPARγ neddylation and CD24 expression in monocytes in a TOLLIP-dependent manner.

### CD24 potentiates inflammation resolution by 4-PBA trained monocytes

CD24 serves as a potent anti-inflammatory mediator and was independently shown to reduce the inflammatory activation of neighboring cells ^48^. We next established a co-culture system to determine whether 4-PBA programmed monocytes effectively propagate inflammation resolution to neighboring cells through CD24. BMMs from B6.SJL mice (CD45.1^+^) were pre-treated with oxLDL to induce inflammatory polarization, and then co-incubated with BMMs from WT B6 or *Cd24*^−/−^ mice (CD45.2^+^) that had been treated with PBS or 4-PBA. After co-culture for 2 days, the cell surface expression of ICAM-1 and CD24 on CD45.1^+^ recipient BMMs was examined by flow cytometry. As shown in Fig. 6a and b, compared with PBS-treated donor monocytes, 4-PBA programmed donor monocytes effectively propagate inflammation resolution to recipient monocytes previously activated by oxLDL as reflected by reduced expression of ICAM-1 and elevated expression of CD24 on recipient monocytes. Importantly, 4-PBA-treated CD24-deficient monocytes failed to impact the expression of ICAM-1 nor CD24 on neighboring recipient monocytes (Figure 6A-B), suggesting that CD24 is required for the propagation of inflammation resolution.

**Figure 6.**
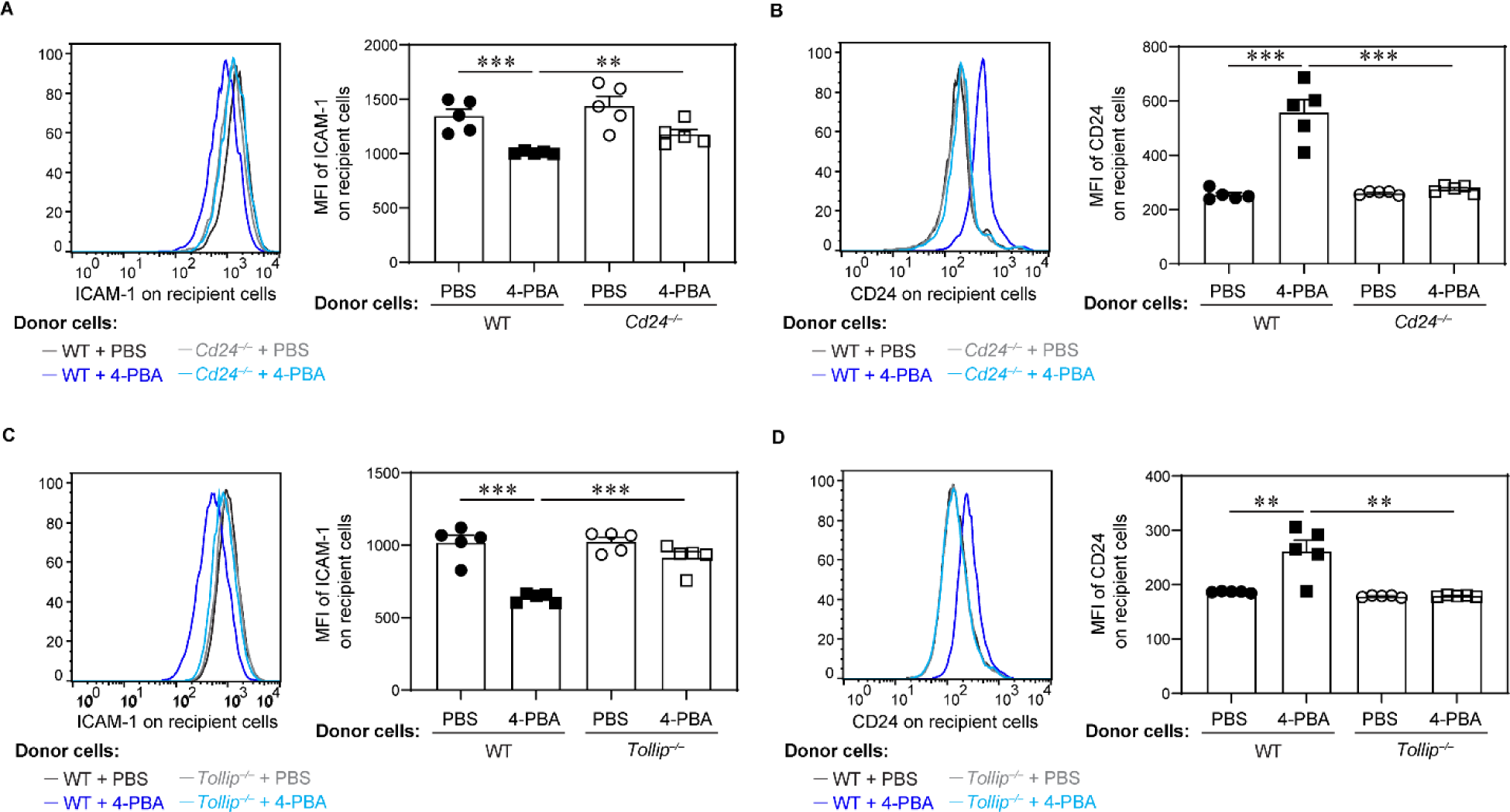
4-PBA primed monocytes propagate resolving nature to neighboring monocytes through CD24 and Tollip. **A-B,** BMMs from WT C57 BL/6 mice and *Cd24*^−/−^ mice, which are both CD45.2^+^, were cultured with M-CSF (10 ng/mL) in the presence of 4-PBA (1 mM) or PBS for 5 days. Recipient BMMs prepared from B6 SJL mice, which are CD45.1^+^, were treated with oxLDL (10 µg/mL) for 3 days and then co-cultured with CD45.2^+^ donor cells for 2 days. Surface expressions of ICAM-1 (**A**) and CD24 (**B**) on CD45.1^+^ recipient BMMs were examined by flow cytometry. **C-D,** BMMs from WT C57 BL/6 mice and *Tollip*^−/−^ mice, which are both CD45.2^+^, were cultured with M-CSF (10 ng/mL) in the presence of 4-PBA (1 mM) or PBS for 5 days. Recipient BMMs prepared from B6 SJL mice, which are CD45.1^+^, were treated with oxLDL (10 µg/mL) for 3 days and then co-cultured with CD45.2^+^ donor cells for 2 days. Surface expression of ICAM-1 (**C**) and CD24 (**D**) on CD45.1^+^ recipient BMMs was examined by flow cytometry. Data are representative of two independent experiments (n = 5 for each group). Error bars represent means ± SEM. \*\**P* < 0.01, \*\*\**P* < 0.001; one-way ANOVA.

Given that we observed TOLLIP is required for PPARγ neddylation and CD24 induction by 4-PBA, we then tested whether *Tollip* deletion may similarly abolish the inflammatory resolution by 4-PBA. Indeed, neither expression of ICAM-1 nor CD24 on neighboring recipient monocytes was significantly impacted by *Tollip*-deficient monocytes trained with 4-PBA (Figure 6C-D).

### 4-PBA trained monocytes effectively reduce atherosclerosis pathogenesis

Although systemic administration of 4-PBA can alleviate atherosclerosis progression, it may also interfere the normal differentiation of monocytes to macrophages, subsequently leading to a decreased number of resident macrophages across various tissues. To enhance the therapeutic specificity of 4-PBA, it is critical to refine its efficacy for targeted purposes. Having defined the phenotypic and mechanistic aspects of resolving monocytes trained by 4-PBA, we then tested whether 4-PBA trained monocytes are sufficient to render atheroprotection when transfused into recipient experimental animals. Both male and female *ApoE^−/−^* mice were subjected to HFD for 4 weeks and then transfused weekly for an additional 4 weeks with BMMs primed with either PBS or 4-PBA. The recipient mice were supplemented with HFD during this process, allowing the development of atherosclerosis. We observed that both male and female recipient mice transfused with 4-PBA–programmed monocytes exhibited a significant reduction in plaque sizes and decreased plaque lipid content (Figure 7A-B and Figure S7A-B), as well as dramatically higher plaque collagen content relative to mice transfused with control monocytes (Figure 7C and Figure S7C). Moreover, mice transfused with 4-PBA–programmed monocytes had significantly lower plasma level of CCL5 (Figure 7D). Given that our *in vitro* data demonstrated 4-PBA-primed monocytes effectively propagate anti-inflammatory activity to neighboring monocytes (Figure 6), we hypothesized that injection of 4-PBA-primed monocytes would induce a similar phenotypic change in monocytes *in vivo*. To test this, we labeled transfused monocytes with carboxyfluorescein succinimidyl ester (CFSE) before injection and then analyzed the surface expression of ICAM-1 and CD24 on CFSE-negative monocytes in recipient mice. We observed elevated CD24 expression on host monocytes in the aorta and bone marrow following the adoptive transfer of 4-PBA-primed monocytes when compared to those receiving PBS-primed monocytes (Figure 7E-F). Injection of 4-PBA-primed monocytes significantly reduced ICAM-1 expression on host monocytes in the aorta but not in the bone marrow (Figure 7E-F), indicating that aortic monocytes of atherosclerotic mice became less adhesive after receiving 4-PBA-programmed monocytes. These data collectively highlight the efficacy of adoptive transfer of 4-PBA– programmed monocytes in mitigating the progression of atherosclerosis. These results also indicate a promising approach of immune cell therapy through utilizing resolving monocytes for the treatment of atherosclerosis.

**Figure 7.**
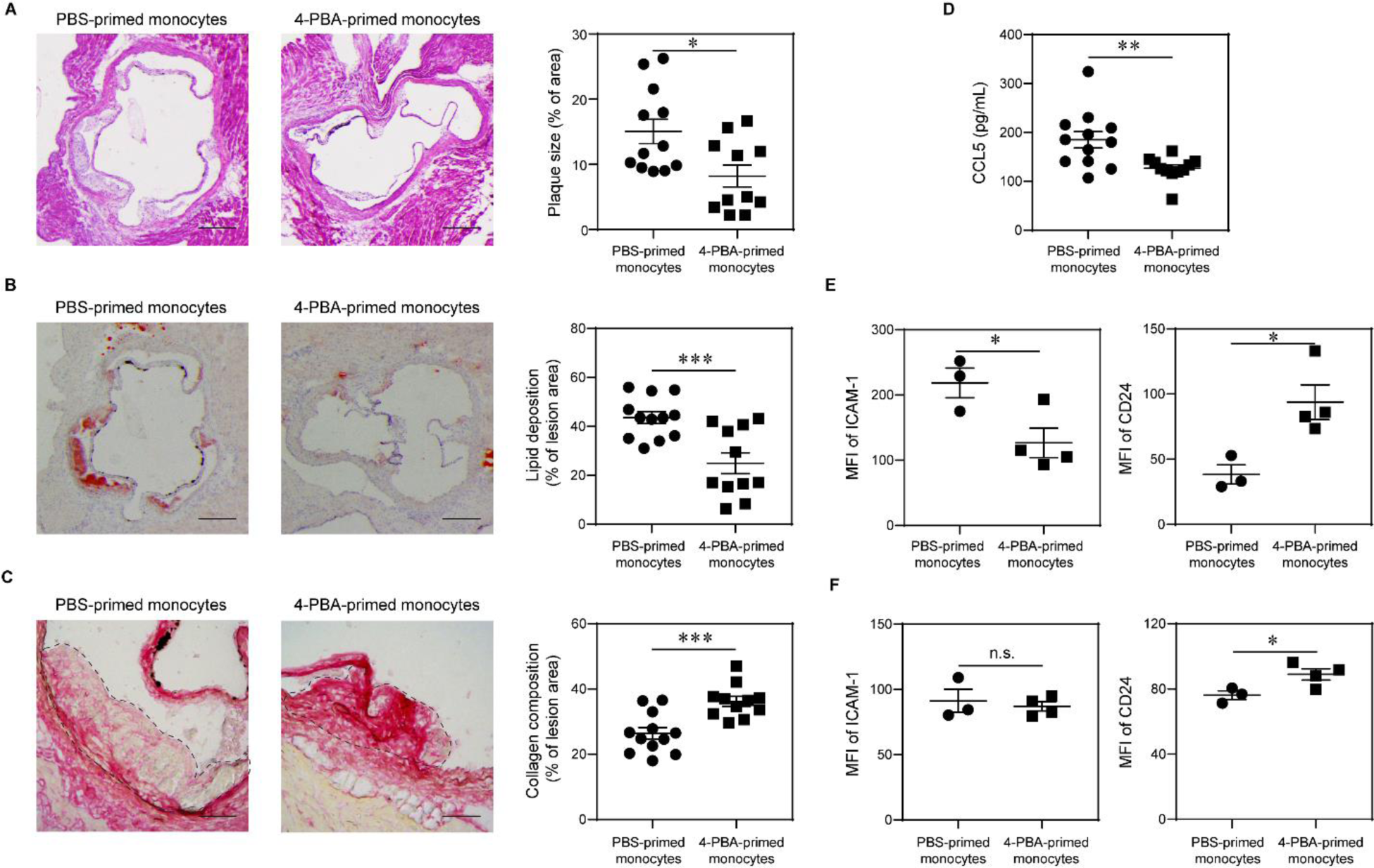
Adoptive transfer of monocytes polarized by 4-PBA alleviates atherosclerosis. **A-D**, Male *ApoE*^−/−^ mice, serving as recipients, were fed with HFD for 4 weeks. BMMs from *ApoE*^−/−^ mice were treated with PBS or 4-PBA (1 mM) for 5 days. PBS-or 4-PBA-polarized monocytes (3 × 10^6^ cells per mouse) were then adoptively transferred by intravenous injection to HFD-fed *ApoE*−/− mice once a week for 4 weeks. Tissues were harvested 1 week after the last monocyte transfer. **A,** Representative images of H&E-stained atherosclerotic lesions and quantification of plaque size demonstrated as the percentage of lesion area within aortic root area. Scale bars, 300 μm. **B,** Representative images of Oil Red O–stained atherosclerotic plaques and quantification of lipid deposition within lesion area. Scale bars, 300 μm. **C,** Representative images of Picrosirius red–stained atherosclerotic plaques and quantification of collagen content within lesion area. Scale bars, 100 μm. **D,** Detection of CCL5 level in the plasma by ELISA. **E-F,** PBS-or 4-PBA-polarized monocytes were labeled with CFSE immediately before adoptive transfer, and tissues were harvested 1 week after the last monocyte transfer. Surface expressions of ICAM-1 and CD24 on host CFSE^−^ CD11b^+^ Ly6G^−^ Ly6C^hi^ monocytes in the aorta (**E**) and BM (**F**) were determined by flow cytometry. Data are representative of two independent experiments (n = 12 for PBS-primed monocytes group and n = 11 for 4-PBA-primed monocytes group in **A-D**; n = 3 for PBS-primed monocytes group and n = 4 for 4-PBA-primed monocytes group in **E** and **F**). Error bars represent means ± SEM. \**P* < 0.05, \*\**P* < 0.01, \*\*\**P* < 0.001, n.s. no statistical significance; Student’s 2-tailed t test.

## Discussion

Our work identified a robust methodology for reprogramming resolving monocytes capable of effectively propagating inflammation resolution and treating atherosclerosis. Monocytes arrested by 4-PBA persist in a less-differentiated, pro-resolving state without further differentiation into the mature and/or inflammatory macrophages typically involved in exacerbating atherosclerosis. 4-PBA trained monocytes exhibit reduced levels of adhesion molecule ICAM-1, immune cell chemokine CCL5, and mature macrophage marker F4/80 while also possessing drastically elevated levels of pro-resolving mediators such as CD24. 4-PBA programmed monocytes can potently propagate inflammation resolution through CD24-mediated inter-cellular communication with neighboring cells. Mechanistically, we demonstrated that 4-PBA effectively reduces monocyte inflammatory signaling by reducing TRAM-mediated cellular stress, as indicated by reduced peroxisome assembly of mTOR and SYK activation. Concurrently, 4-PBA robustly promotes to the expression of anti-inflammatory CD24 by facilitating TOLLIP-mediated PPARγ-neddylation and activation.

Our study outlined a novel immune cell-based therapeutic alternative for the future treatment of atherosclerosis. Previously, 4-PBA was shown to be effective in treating experimental atherosclerosis ^19–21^. However, systemic administration of chemical compounds such as 4-PBA introduces a plethora of side effects, potentially altering host defense, cellular development, and other vital tissue functions. Our *ex vivo* characterization demonstrated that monocytes treated with 4-PBA are arrested at a less-mature monocytic state, which suggests that long-term systemic injection of 4-PBA would likely impact systemic macrophage development and host anti-microbial defense. By contrast, administration of 4-PBA-reprogrammed monocytes would enable clinicians to selectively harness their anti-inflammatory and therapeutic potential while alleviating the side effects of systemic 4-PBA application.

Through integrated scRNA-seq and relevant functional analyses, our current work not only presents a systematic characterization of 4-PBA programmed resolving monocytes, but also reveals key mechanistic insights into monocyte reprogramming dynamics. We determined that 4-PBA reprograms resolving monocytes by reducing peroxisome-mediated mTOR and SYK signaling circuitry. Our findings validate previous reports that mTOR inhibitors such as rapamycin serve as effective agents in reducing atherosclerosis ^49^. Nanoparticle mediated delivery of mTOR inhibitors were reported to be effective in treating experimental atherosclerosis ^50,51^. Our mechanistic studies further defined the TRAM adaptor as a membrane-associated stress sensor of inflamed monocytes and demonstrated that 4-PBA can effectively reduce TRAM-mediated membrane stress. TRAM is one of the few innate signaling adaptors with covalently-associated lipid motifs allowing them to be anchored onto the membrane lipid leaflet ^52,53^. Lipid-modifications such as palmitoylation or myristoylation not only serve to anchor signaling molecules to the cell membrane ^54–56^, but also can facilitate the sensing of membrane stress signals independent of cell surface receptors ^57,58^. Previous studies revealed that increased membrane stress and/or rigidity can facilitate the assembly of lipid raft, where lipid-conjugated protein adaptors such as TRAM can dock and undergo activation ^59–62^. Independent reports demonstrate that cholesterol and/or oxLDL can generically increase membrane rigidity and lipid raft formation ^39–41^. Complementing these studies, our current work revealed that oxLDL indeed can cause membrane clustering of the TRAM adaptor in activated monocytes. Our functional data further validated that TRAM may serve as a general membrane stress sensor for monocyte activation given that the induction of ICAM-1 and CCL5 by oxLDL is ablated in TRAM-deficient monocytes. Our data can also reconciles previous independent findings that implicate TRAM as a key signaling adaptor for oxidized phospholipid-induced monocyte activation ^63,64^. Our data demonstrating the role of 4-PBA in alleviating TRAM clustering and reducing monocyte inflammatory polarization suggest that additional compounds that relieve membrane stress may be similarly effective in dampening monocyte low-grade inflammatory memory.

Our data reveal the key role of CD24 plays in 4-PBA trained monocytes during the propagation of anti-inflammatory resolution. This is consistent with emerging studies reporting the beneficial roles of CD24 in reducing tissue inflammation ^30,65–67^. CD24 has been shown to inhibit inflammatory activation in neighboring cells through ligating its cognate inhibitory receptor Siglec-10 ^48,65^. A recombinant CD24-immunoglobulin protein has been shown to exert anti-inflammatory effects for the treatment of chronic inflammatory diseases such as diabetes ^30,65^. Our current study outlines a novel approach for reprogramming monocytes into a homeostatic resolution state with enhanced cellular expression of CD24, thereby providing a targeted and effective immune cell-based anti-inflammatory therapy capable of treating atherosclerosis and perhaps other chronic inflammatory conditions.

Taken together, our study demonstrates the feasibility and effectiveness of 4-PBA trained monocytes for promoting homeostatic resolution *in vitro* and *in vivo* by programming monocytes into a less-differentiated, non-inflammatory, and CD24-expressing homeostatic resolving state. We further demonstrate the presence as well as the expansion of CD24-expressing intermediate monocytes by 4-PBA treatment *ex vivo*, suggesting the feasibility of generating resolving primary human monocytes. Further refinement of innate monocyte-based approaches harbors tremendous promise for generating immune cell-based precision therapies for atherosclerosis.

## Acknowledgements

The authors would like to acknowledge the assistance of Li lab members for the animal breeding and care, as well as technical assistance.

---

**Figure S1.**
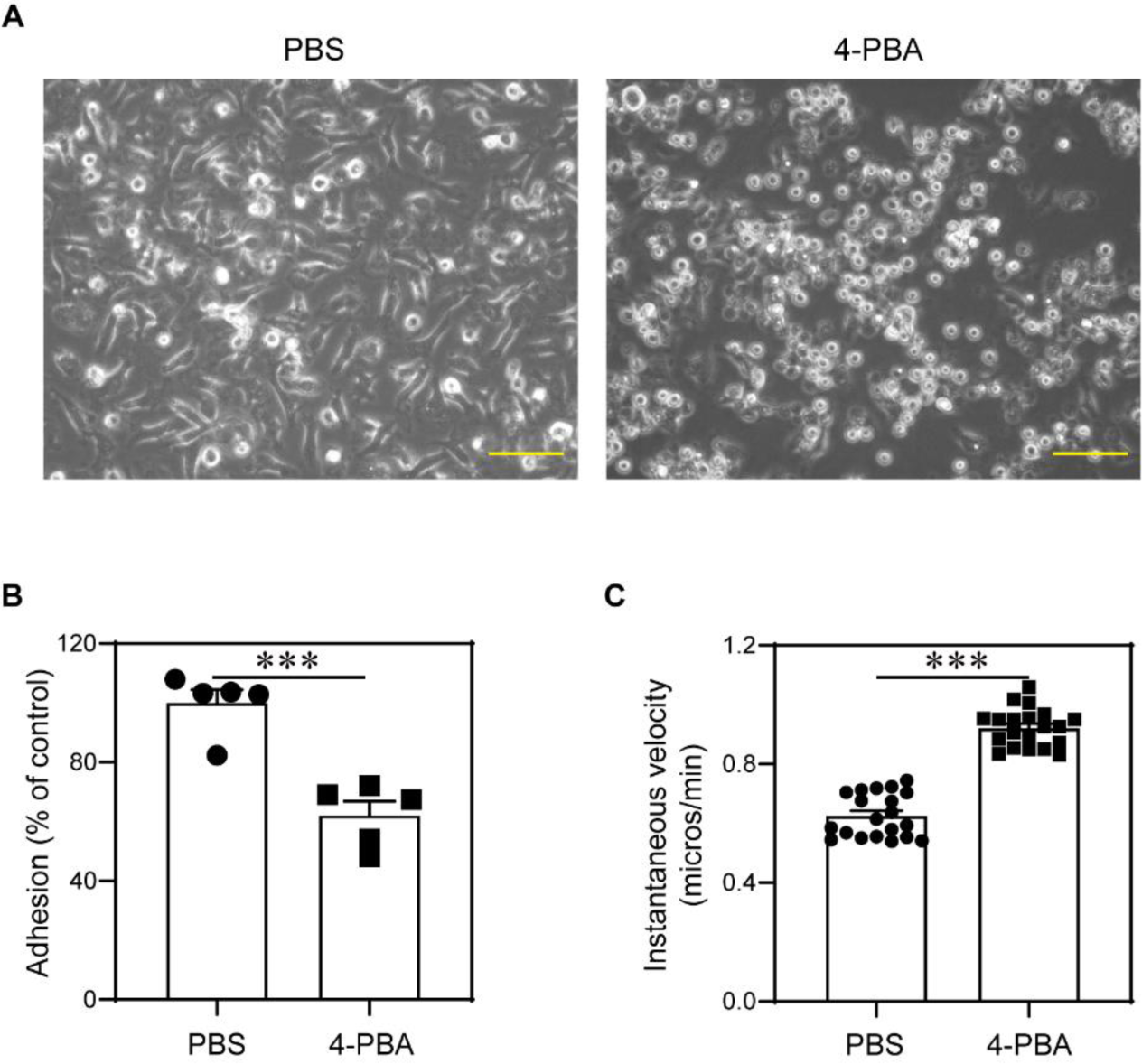
Treatment with 4-PBA induces morphological and behavioral changes in monocytes. BMMs from WT C57 BL/6 mice were cultured with M-CSF (10 ng/mL) in the presence 4-PBA (1 mM) or PBS for 5 days. **A,** Morphology of monocytes was observed under a conventional bright field microscope. Scale bars, 50 μm. Adhesiveness (**B**) and migratory capacity (**C**) of monocytes were evaluated. Data are representative of three independent experiments (n = 5 for each group in **B**; n = 20 for each group in **C**). Error bars represent means ± SEM. \*\*\**P* < 0.001; Student’s 2-tailed t test.

**Figure S2.**
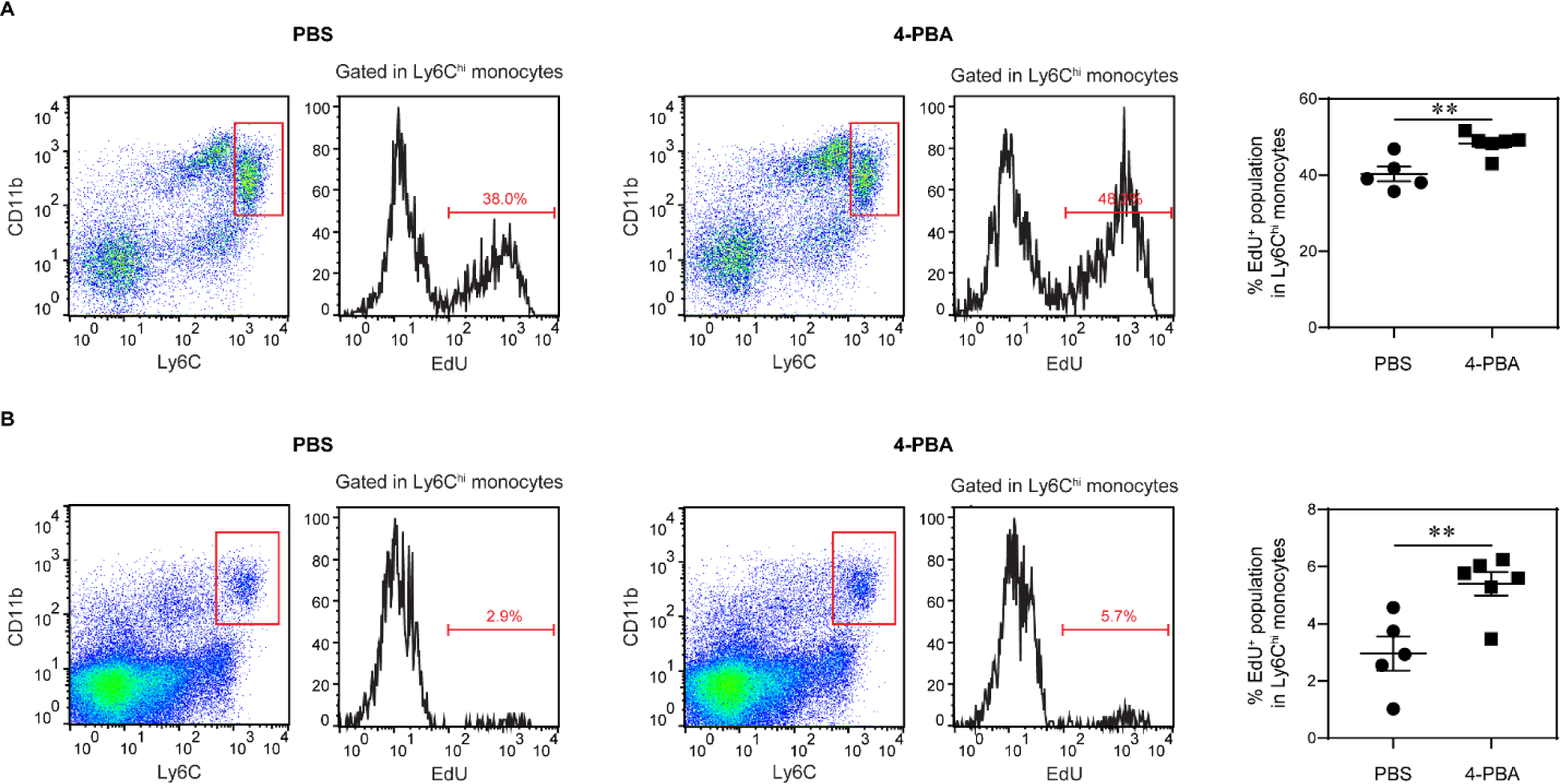
Injection of 4-PBA promotes monocyte proliferation in atherosclerotic mice. Male *ApoE*^−/−^ mice were fed with HFD for 4 weeks and intraperitoneally injected with 4-PBA (5 mg/kg body weight) or PBS every 3 days for additional 4 weeks. EdU (50 mg/kg body weight) was intraperitoneally injected to the mice 4 hours before euthanasia. EdU^+^ proliferating population in CD11b^+^ Ly6G^−^ Ly6C^hi^ monocytes in the BM (**A**) and spleen (**B**) was detected by flow cytometry. Data are representative of two independent experiments (n = 5 for PBS group and n = 6 for 4-PBA group). Error bars represent means ± SEM. \*\**P* < 0.01; Student’s 2-tailed t test.

**Figure S3.**
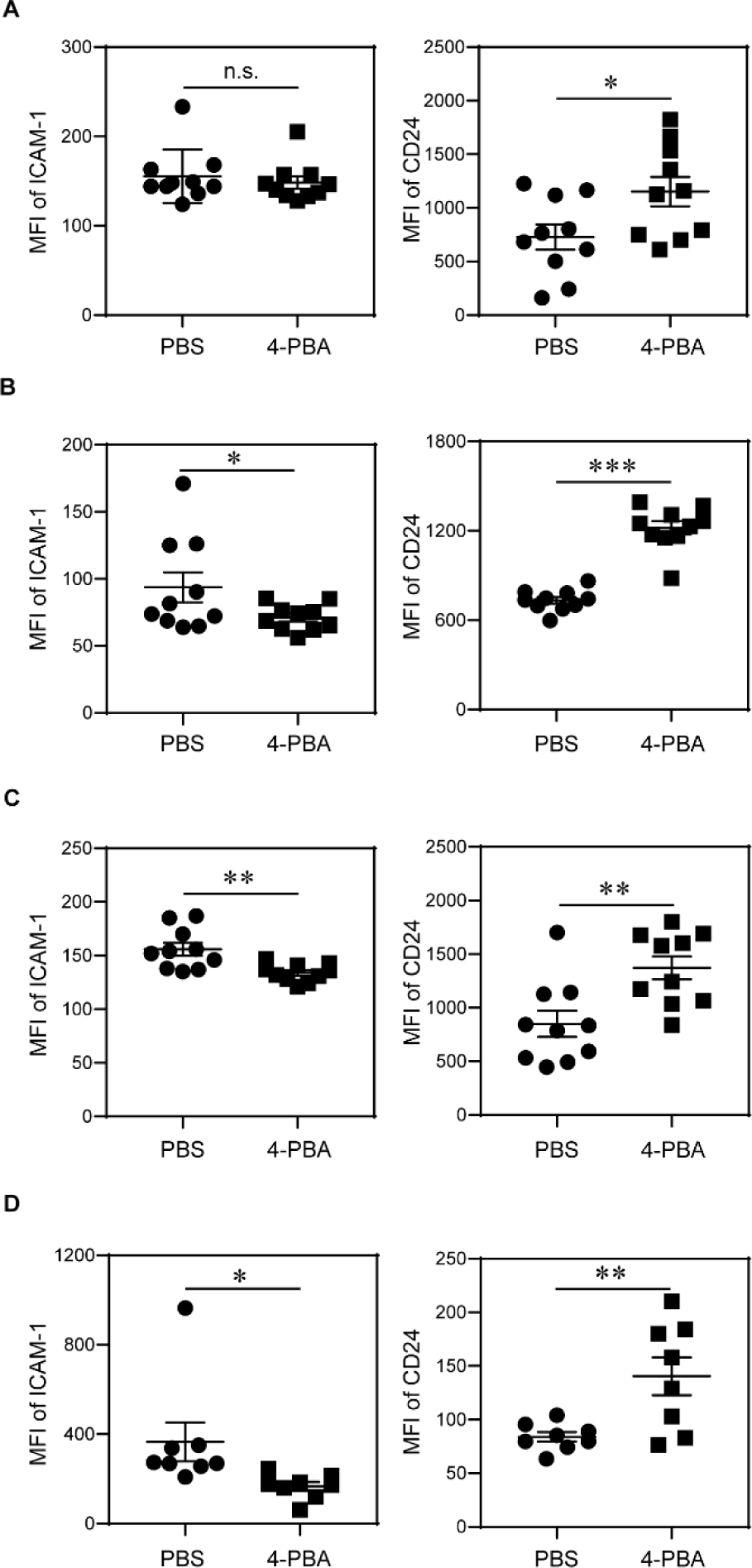
Administration of 4-PBA reprograms monocytes in vivo. Male *ApoE*^−/−^ mice were fed with HFD for 4 weeks and intraperitoneally injected with 4-PBA (5 mg/kg body weight) or PBS every 3 days for additional 4 weeks. **A-D,** Surface expressions of ICAM-1 and CD24 on CD11b^+^ Ly6G^−^ Ly6C^low^ monocytes in the peripheral blood (**A**), BM (**B**), spleen (**C**), and aorta (**D**) were examined by flow cytometry. Data are representative of two independent experiments (n = 10 for each group in **A-C**; n = 8 for each group in **D**). Error bars represent means ± SEM. \**P* < 0.05, \*\**P* < 0.01, \*\*\**P* < 0.001, n.s. no statistical significance; Student’s 2-tailed t test.

**Figure S4.**
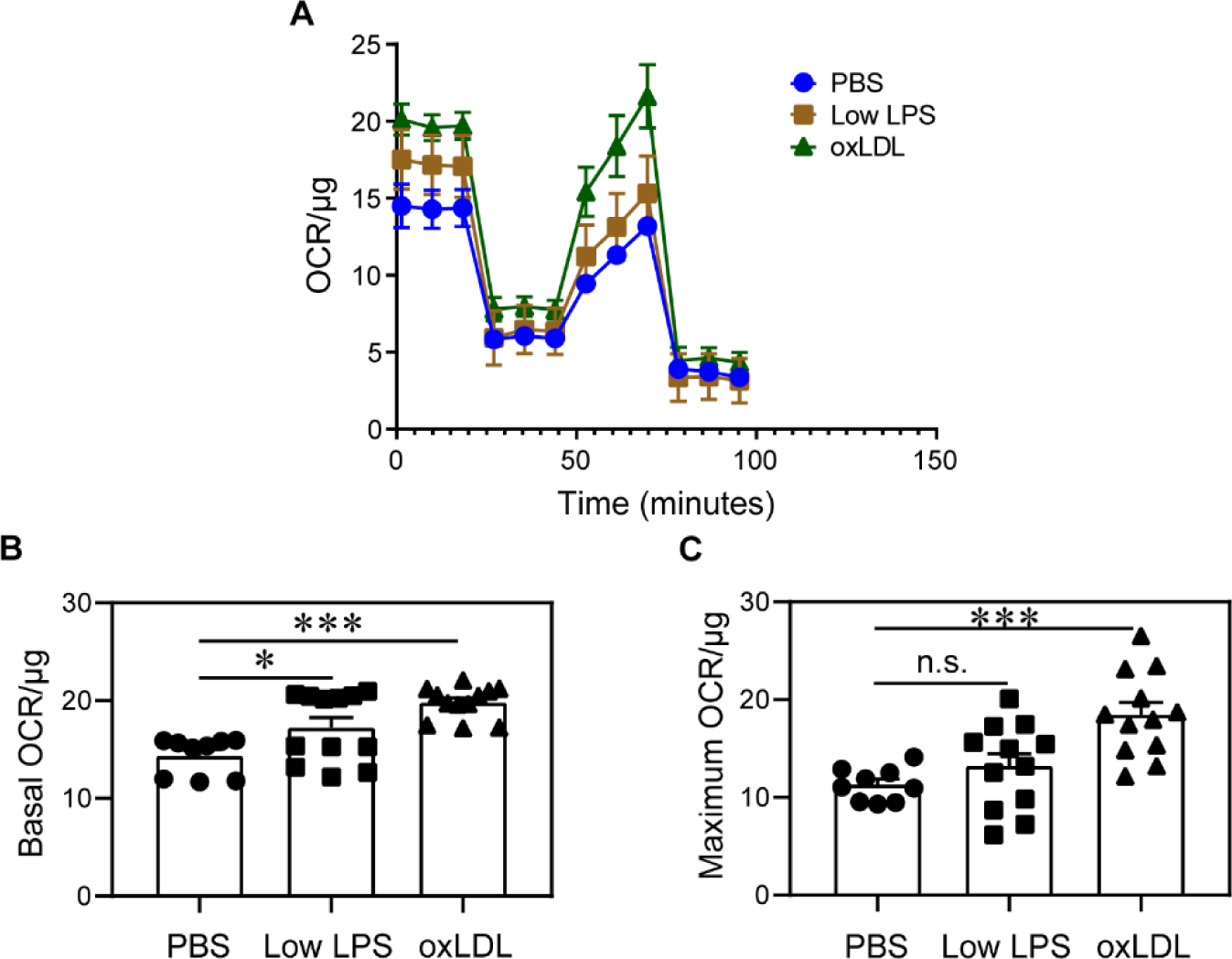
Low dose LPS and oxLDL exert similar effects in enhancing the mitochondrial respiration of monocytes. BMMs from WT C57 BL/6 mice were cultured with M-CSF (10 ng/mL) in the presence of low dose LPS (100 pg/mL), oxLDL (10 µg/mL), or PBS for 5 days. **A,** Oxygen consumption rate (OCR) of each group was determined by Seahorse assay. **B-C,** The basal and maximum OCR/ug levels were quantified. Data are representative of three independent experiments (n = 9 for PBS group, n = 12 for Low LPS group and ox LDL group). Error bars represent means ± SEM. \**P* < 0.05, \*\*\**P* < 0.001, n.s. no statistical significance; one-way ANOVA.

**Figure S5.**
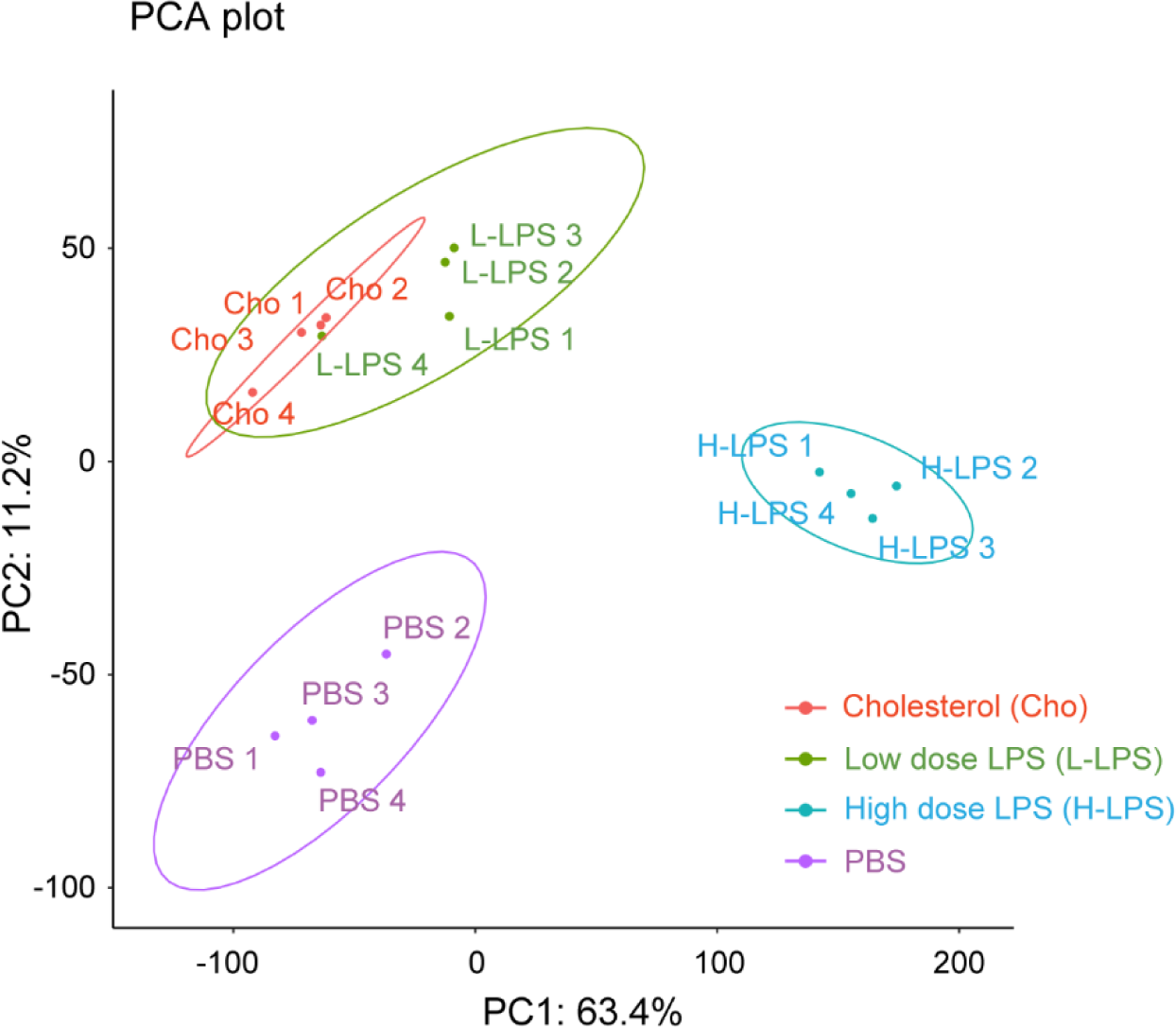
Low dose LPS and cholesterol stimulation elicits similar DNA methylation reprogramming in monocytes. BMMs from WT C57 BL/6 mice were cultured with M-CSF (10 ng/mL) in the presence of low dose LPS (100 pg/mL), high dose LPS (1 µg/mL), cholesterol (10 µg/mL) or PBS for 5 days. DNA methylation profiles were analyzed. Principal component analysis of differentially methylated CpG probes in monocytes was conducted. Ovals indicate the normal distribution for each treatment.

**Figure S6.**
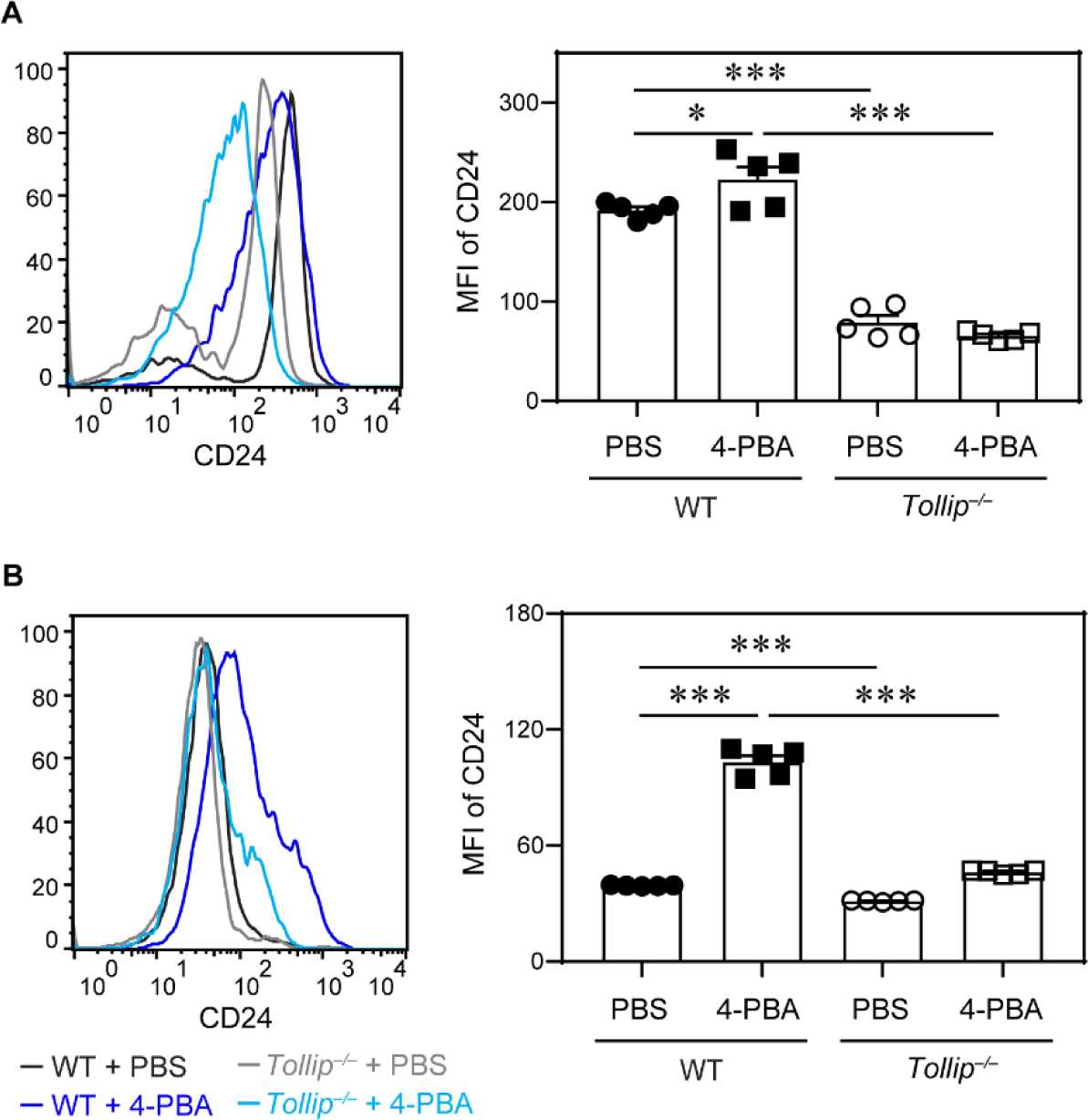
4-PBA promotes the expression of CD24 in a Tollip dependent manner. BMMs from WT C57 BL/6 mice and *Tollip*^−/−^ mice were cultured with M-CSF (10 ng/mL) in the presence of 4-PBA (1 mM) or PBS for 5 days. Surface expression of CD24 on CD11b^+^ Ly6C^low^ monocytes (**A**) and CD11b^+^ Ly6C^−^ monocytes (**B**) was examined by flow cytometry. Data are representative of three independent experiments (n = 5 for each group). Error bars represent means ± SEM. \**P* < 0.05, \*\*\**P* < 0.001; one-way ANOVA.

**Figure S7.**
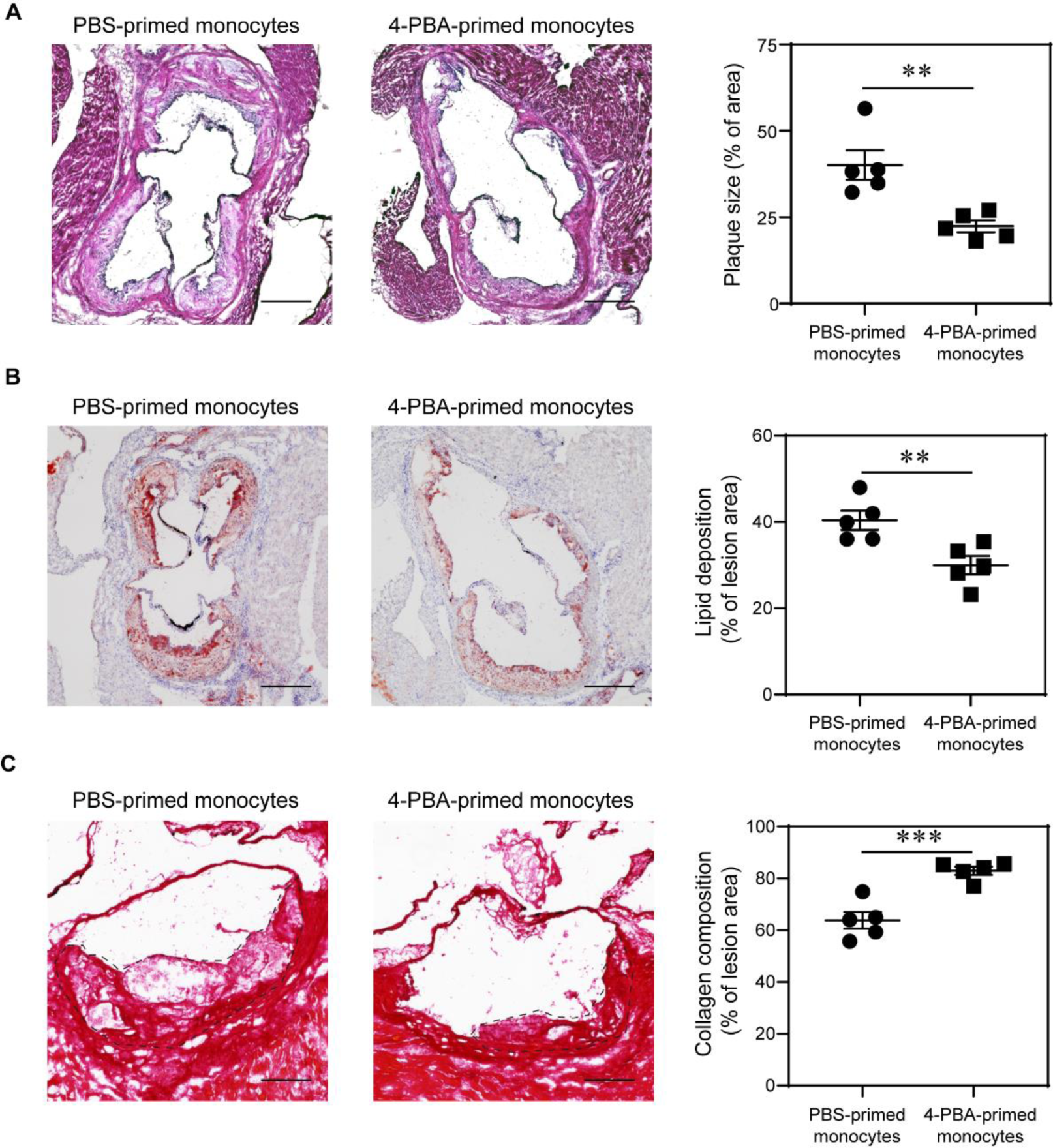
Adoptive transfer of monocytes polarized by 4-PBA alleviates atherosclerosis. Female *ApoE*^−/−^ mice, serving as recipients, were fed with HFD for 4 weeks. BMMs from *ApoE*^−/−^ mice were treated with PBS or 4-PBA (1 mM) for 5 days. PBS-or 4-PBA-polarized monocytes (3 × 10^6^ cells per mouse) were then adoptively transferred by intravenous injection to HFD-fed *ApoE*−/− mice once a week for 4 weeks. Tissues were harvested 1 week after the last monocyte transfer. **A,** Representative images of H&E-stained atherosclerotic lesions and quantification of plaque size demonstrated as the percentage of lesion area within aortic root area. Scale bars, 300 μm. **B,** Representative images of Oil Red O–stained atherosclerotic plaques and quantification of lipid deposition within lesion area. Scale bars, 300 μm. **C,** Representative images of Picrosirius red–stained atherosclerotic plaques and quantification of collagen content within lesion area. Scale bars, 100 μm. Data are representative of two independent experiments (n = 5 for each group). Error bars represent means ± SEM. \*\**P* < 0.01, \*\*\**P* < 0.001; Student’s 2-tailed t test.

## Non-standard Abbreviations and Acronyms

4-PBA: 4-phenylbutyric acid
BMM: bone marrow-derived monocyte
CCL5: Chemokine ligand 5
CFSE: Carboxyfluorescein succinimidyl ester
H&E: Hematoxylin and Eosin
HFD: high fat diet
ICAM-1: Intercellular adhesion molecule 1
M-CSF: Macrophage colony-stimulating factor m
TOR: Mammalian target of rapamycin
NEDD8: Neural precursor cell expressed, developmentally down-regulated 8
OCT: optimal cutting temperature
oxLDL: Oxidized low-density lipoprotein
PPARγ: Peroxisome proliferator-activated receptor γ
scRNA-seq: Single-cell RNA sequencing
SYK: Spleen tyrosine kinase
TOLLIP: Toll-interacting protein
TRAM: Trif-related adapter molecule

